# Network Correlates of Shared Experience in Musical Improvisation

**DOI:** 10.64898/2026.11.25.768844

**Authors:** Alberto Liardi, Madalina I. Sas, Hardik Rajpal, Ionuţ-Flavius Bratu, Fernando E. Rosas, Pedro A.M. Mediano, Henrik J. Jensen

## Abstract

Shared experiences emerge when individuals feel attuned to one another as they coordinate their actions and mental states, constituting a fundamental part of everyday human life. Yet, the neural mechanisms underpinning these phenomena remain poorly understood. Musical improvisation provides an ecologically valid setting for studying shared experiences, as it combines creativity, joint action, and real-time adaptation. Here, we investigate the neural correlates of shared experience during group musical performance using hyperscanning EEG recordings of professional musicians performing in naturalistic settings in front of an audience. Musicians performed each piece under two conditions: a *strict* rendition, focused on precise execution of the score, and a *let-go* version, which encourages improvisation and risk-taking. This approach enables the isolation of neural signatures specifically associated with improvisational and shared states of mind, which we probe using tools from information theory and network science. Our results reveal that, across most pieces and ensembles, *let-go* performances are associated with greater signal complexity and interdependence within and between brains, alongside denser, more cohesive information flow and a more synergistic combination of the musicians’ brain networks. These findings point to more integrated collective neural dynamics during *let-go*, consistent with the emergence of a shared experience, and suggest that neural signatures of conscious states may extend beyond individual brains to interacting systems. Beyond these empirical findings, this work introduces a methodological toolkit combining complexity measures, higher-order information-theoretic metrics, and network-based analyses to investigate inter-brain coordination and collective cognition in naturalistic settings.

## I. INTRODUCTION

From teamwork and artistic collaboration to collective celebration, human life is rich in situations in which individuals experience a heightened sense of intercon-nectedness and shared awareness. Such *shared experiences* (Chung et al. 2024) are characterised by enhanced empathy and a sense of unity (Rennung and Göritz 2016; Tarr et al. 2014), and can foster a collective mindset that seems to transcend individual thought processes (Combs and Kripner 2008; Midgley 2006; Noy et al. 2015; Schmid 2014), even described by some as akin to “mind-reading” (Hart and Di Blasi 2015; Sawyer 2003). Shared experience appears closely tied to a group’s capacity to coordinate or cooperate towards a common goal, such as during artistic collaboration, problem-solving, or group discussions, but also during purely communal activities (Combs and Kripner 2008; Midgley 2006; Noy et al. 2015; Schmid 2014). However, while collective activities are often investigated from a cognitive or sen-sorimotor perspective, their phenomenological counter-part – the shared experience that accompanies successful coordination – remains understudied, partly because it is unclear which properties of neural activity should be measured to capture it.

Neuroscience of consciousness offers a principled approach to address this question by identifying properties of neural activity that track conscious states. Three signatures have been reported as particularly useful for tracking the richness of a conscious state: *complexity, integration*, and *synergy*. A decrease in signal complexity (Casali et al. 2013; Sarasso et al. 2015, 2021) and long-range integration of differentiated activity (Demertzi et al. 2019; Luppi et al. 2019; Tononi 2004; Tononi and Edelman 1998; Tononi et al. 1994) have been found to reliably predict loss of consciousness in dreamless sleep, general anaesthesia, and disorders of consciousness. Similarly, a reduced level of consciousness also correlates with weaker synergistic emergent dynamics, in which information is carried only by the joint state of multiple components (Luppi et al. 2023). Moreover, these signatures also vary in a graded way among conscious states: complexity rises above wakeful levels under psychedelics (Mediano et al. 2024; Schartner et al. 2017) and fluctuates with cognitive engagement during ordinary wakefulness (Mediano et al. 2021); integration increases under psychedelics and correlates with ego dissolution (Tagliazucchi et al. 2016), and synergy is most prominent in regions supporting higher cognitive function (Luppi et al. 2022, 2024b).

The effectiveness of these neural correlates of conscious experience suggests an approach to studying shared experience: if shared experience corresponds to a genuinely joint state (rather than to separate individual experiences that merely co-occur), these signatures should not only be enhanced within each individual’s brain, but also extend to the joint system formed by all participants together. Importantly, testing this hypothesis requires an appropriate experimental setting in which shared experience can be elicited and contrasted against a baseline.

Musical performances are ideal for examining both individual and interpersonal phenomenology, as they broadly engage the spectrum of consciousness: emotional expression (Juslin and Sloboda 2010), flow (Zielke et al. 2023), absorption (Cardona et al. 2022), mind-wandering (Palhares et al. 2024), and even psychedelic-like states (Kaelen et al. 2018; Kathofer et al. 2025). Furthermore, group music performance requires both shared phenomenology and joint action (Keller 2008): musicians coordinate movements (Chang et al. 2019; Walton et al.

2015), distribute attention, and synchronise sensorimotor patterns across brains (Keller et al. 2014). In particular, collective musical improvisation fuses flow, creativity, and risk-taking into a distinct state of mind characterised by a high neural entropy (Dolan et al. 2018), making it a natural candidate for a richer conscious state. Group music performance is, thus, an ideal setting for hyperscanning (Acquadro et al. 2016), the study of synchronised brain activity across interacting participants (Czeszumski et al. 2020; Zamm et al. 2024), where it has been associated with real-time adaptive inter-brain coupling expressed as synchrony (Sänger et al. 2012; Zamm et al. 2021) or information flow (Sänger et al. 2013). However, existing hyperscanning studies of improvisation fall short of the requirements of shared phenomenology and joint action, often relying on constrained tasks, restricted lab setups, and heterogeneous analysis pipelines (Acquadro et al. 2016; Shamay-Tsoory and Mendelsohn 2019; Zimmermann et al. 2024).

An emerging research paradigm in classical music improvisation (Dolan et al. 2018, 2013) addresses these issues by using mobile sensors on professional musicians who play entire movements in a naturalistic, real-life performance space with real instruments and an audience. This practice also offers a unique baseline: each contrast compares a “*strict* “ performance, focused on reproducing the score accurately and expressively, against a “*let-go*” performance, where risk-taking is encouraged, and ensemble partners may spontaneously deviate from the score (Dolan et al. 2018). This comparison isolates the specific effects of improvisation, both in the brain and in music performance parameters.

**TAB I:** Summary of datasets studied in this work. “Channels” refers to the number of EEG electrodes per subject.

| **Name** | **Composer** | **Instruments** | **Channels** | **Sampling freq.** | **Freq. range** | **Dataset** |
|---|---|---|---|---|---|---|
| Piece 1 | Beethoven | Piano, Cello | 16 | 250 Hz | 1–125 Hz | New |
| Piece 2 | Schumann | Piano, Cello | 16 | 250 Hz | 1–125 Hz | New |
| Piece 3 | Schubert | Flute, Piano, Singer | 19 | 250 Hz | 2–40 Hz | Dolan et al. (2018) |
| Piece 4 | Ravel | Flute, Harp, Viola | 10 | 100 Hz | 1–50 Hz | Dolan et al. (2013) |
| Piece 5 | Improvisation | Flute, Harp, Viola | 10 | 100 Hz | 1–50 Hz | Dolan et al. (2013) |

On the analytical side, even under a robust experimental setup, symmetric synchrony measures struggle to disentangle genuine interaction-driven coupling from correlations induced by shared stimuli or coordinated motor timing (Markus and Shamay-Tsoory 2024). Beyond this, the synchronyor correlation-based measures used in most existing studies are inherently pairwise: they quantify dependencies between two signals at a time and therefore cannot capture higher-order interactions, i.e. dependencies that emerge only from the joint activity of three or more components, which are increasingly recognised as key to characterising cognition and conscious states (Liardi et al. 2025; Luppi et al. 2023, 2022, 2024b). Additionally, they quantify the overall strength of inter-dependence without distinguishing between qualitatively different forms of interaction, such as redundancy and synergy (Mediano et al. 2022; Rosas et al. 2026). This lack of an interpretable framework for characterising both the structure and nature of interactions makes it difficult to compare findings across studies, as well as to relate observed inter-brain dependencies to the signatures of conscious states.

Information theory and network science provide a principled framework for addressing these limitations, enabling the characterisation of directed and higher-order dependencies within and across brains, while distinguishing different modes of information sharing and retaining a clear operational interpretation (Luppi et al. 2024b; Mediano et al. 2022; Rosas et al. 2019). Moreover, since the measures of complexity, integration, and synergy used in consciousness science are themselves formulated in information-theoretic terms, they can be naturally extended from single brains to the joint system of interacting brains.

In this paper, we therefore combine the *strict* /*let-go* paradigm with tools from information theory and network science to characterise the neural correlates of shared experience during musical group improvisation. Specifically, we test whether complexity, integration, and synergy are enhanced during *let-go* performances, and whether this enhancement extends across brains. To this end, we introduce a new hyperscanning EEG dataset from a professional piano–cello duo performing two classical pieces in both *strict* and *let-go* modes, and complement it with analyses of two previously published datasets of classical chamber trios (Dolan et al. 2018, 2013). We first replicate the increase in signal complexity during *let-go* reported in earlier work (Dolan et al. 2018), showing that this effect is concentrated in the *γ*

band. We then show that *let-go* performances are characterised by stronger statistical interdependence (integration) both within and between brains, denser and more cohesive information flow, and a more synergistic combination of the musicians’ brain networks. Together, these findings provide a quantitative link between improvisational states and collective neural dynamics, while establishing a methodological framework for investigating shared experience in collective behaviour beyond musical performance.

## II. RESULTS

To ensure generalisability and robustness, we analysed EEG data from three sources of improvised classical chamber music (Tab. I): a novel experiment with a piano–cello duo (Pieces 1–2, see Sec. IV F), and two previously published experiments with classical trios (Pieces 3–5) (Dolan et al. 2018, 2013), which we re-analyse here to test whether our findings generalise across ensembles and recording setups. Each piece in every dataset was performed in both *strict* and *let-go* modes (Dolan et al. 2018, 2013), enabling a direct comparison between non-improvisational and improvisational states. In total, the data comprise eight professional musicians and five pieces, recorded with 10 to 19 EEG channels per musician and sampling frequencies between 100 and 250 Hz (see Tab. I for details). All recordings were filtered and cleaned of bad channels, epochs, and artefacts, then z-scored and segmented into epochs of 1,000 samples, corresponding to 4 s for Pieces 1–3 and 10 s for Pieces 4–5. Further details on the data and preprocessing steps can be found in Sec. IV F.

Our analyses are organised from individual to collective levels of description, each probing a different facet of shared experience (Fig. 1). We start from the signal complexity of each musician’s brain, a well-established marker of the richness of conscious states. We then move to the multivariate interdependencies within and across brains, which quantify how strongly neural activity becomes integrated during performance. Next, we construct information-flow networks spanning all musicians, investigating the balance between segregation and integration of the joint system. Finally, we decompose these networks to quantify the extent to which the brain networks of the musicians complement each other from an information-communication perspective. Together, this analysis pipeline allows us to assess whether the emergence of shared experience is accompanied by increased neural interdependence between the musicians, and whether signatures of conscious states extend from individual brains to the joint system.

**FIG 1:**
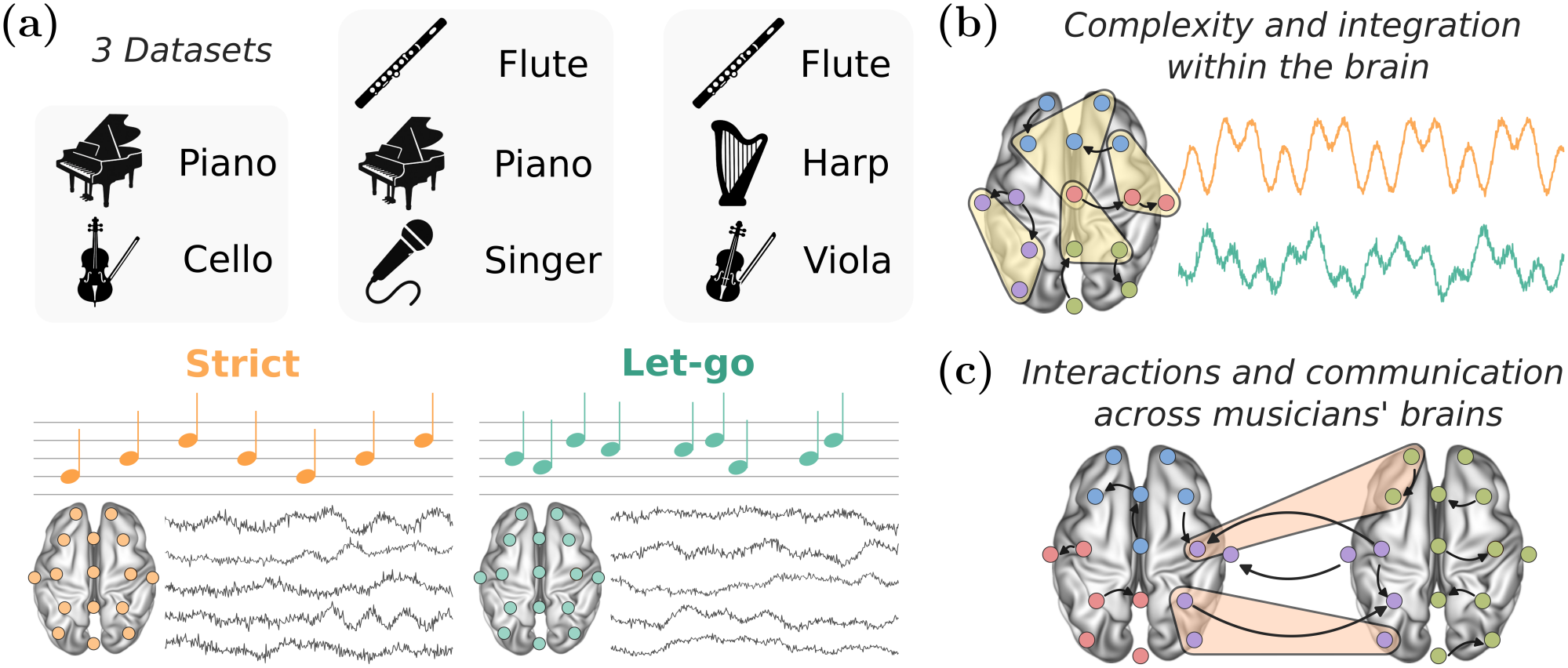
Overview of the datasets and analyses. (a) We analyse hyperscanning EEG recordings of professional musicians from three classical ensembles: a Piano–Cello duo (Pieces 1–2, new dataset), a Flute–Piano–Singer trio (Piece 3), and a Flute–Harp–Viola trio (Pieces 4–5; see Tab. I). Each piece was performed twice, in *strict* and *let-go* modes. (b) Within each brain, we compare the complexity of the neural signals between the two modes (Complexity via State-space Entropy Rate, CSER), and quantify the interdependence among groups of electrodes (S- and O-information). (c) Across brains, we quantify the interdependence among electrodes of different musicians, build information-flow networks spanning the brains of all musicians (Partial Mutual Information from Mixed Embedding, PMIME), and quantify how the brain networks of different musicians complement each other (Partial Network Decomposition, PND).

### A. Signal complexity

Signal complexity is among the most widely used markers of neural dynamics across states of consciousness (Akar et al. 2015; Bachmann et al. 2015; Burioka et al. 2005; Imperatori et al. 2021; Lau et al. 2022; Li et al. 2008; Mediano et al. 2023, 2024; Rajpal et al. 2022; Sarasso et al. 2021; Schartner et al. 2015, 2017; Zhang et al. 2001). According to the entropic brain hypothesis, the richness of conscious experience is reflected in the entropy of spontaneous brain activity (Carhart-Harris 2018; Carhart-Harris et al. 2014), making complexity measures natural candidates to track changes in the quality of the musicians’ experience across performance modes. Within improvisational performance, increased signal complexity has previously been reported in *let-go* compared to *strict* modes (Dolan et al. 2018). Here we quantified complexity using Complexity via State-space Entropy Rate (CSER) (Mediano et al. 2023), a recently introduced estimator which operates directly on continuous data and allows a frequency-resolved decomposition of complexity (see Sec. IV), offering advantages over classical estimators used in the existing literature.

Across the three experiments, results indicate that overall CSER is higher during *let-go* than *strict* performances (Fig. 2a), with significant increases for five of the 13 musician–piece combinations, spanning all three ensembles (e.g. piano in Piece 1: *t*(15) = 9.80, *p*_FDR_ *<* 10^*™*6^, *d*_*z*_ = 2.45; viola in Piece 5: *t*(9) = 7.35, *p*_FDR_ *<* 10^*™*3^, *d*_*z*_ = 2.33). Even in cases that do not reach statistical significance – likely due to the short duration of the musical excerpts – the observed distributions still suggest a consistent trend towards increased signal complexity in the *let-go* condition, with 12 of the 13 combinations showing a positive mean difference. Accordingly, when CSER is averaged across channels, and the 13 musician–piece combinations are taken as samples, the increase is significant across the whole dataset (*t*(12) = 4.61, *p* = 3*×*10^*™*4^, *d*_*z*_ = 1.28; Tab. S1). Taken together, these findings suggest that each musician’s brain shifts towards a richer and more differentiated activity during *let-go*, replicating the results of Dolan et al. (2018) and providing a foundation for further analysis.

**FIG 2:**
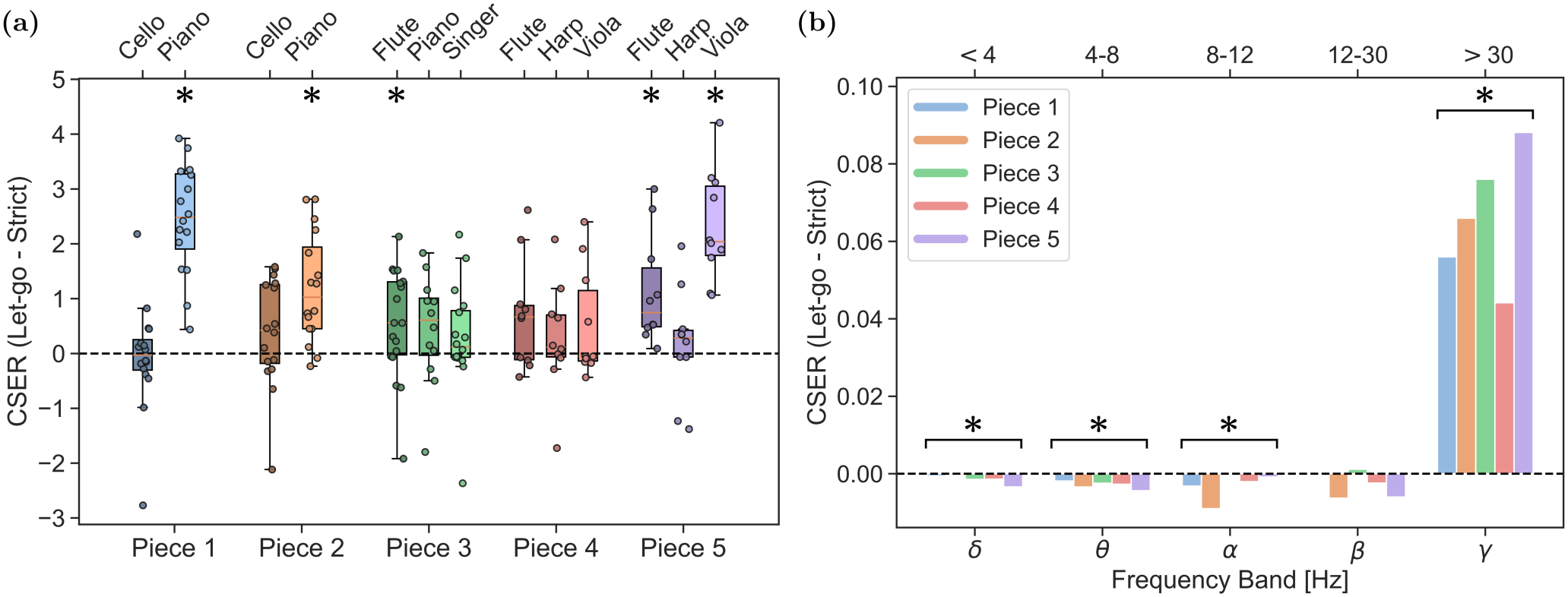
Complexity of neural activity increases during musical improvisation. **(a)** *Let-go*–*Strict* difference in broadband CSER for each piece and musician, with each point representing an EEG electrode. For each musician, the observed differences are normalised by their standard deviation across electrodes so that each distribution has unit variance. Thus, the mean difference can be interpreted as a standardised effect size (Cohen’s *d*_*z*_, see Sec. IV G). *Let-go*–*Strict* CSER decomposed into frequency bands for each piece and averaged across musicians. The *γ* band comprises frequencies above 30 Hz, with an upper limit that depends on the dataset (Tab. I). Stars mark significant differences (one-sample t-test on the paired differences, false discovery rate (FDR) corrected *p <* 0.05; see Sec. IV G and Tab. S1); in (b), each band is tested across all 13 musician–piece combinations, and the corresponding stars are shown above brackets spanning the five pieces.

Leveraging the flexibility of CSER, we then performed a spectral decomposition of signal complexity to examine the contributions of different frequency bands (Fig. 2b). Across all musicians and pieces, the *γ* band shows a significant increase in *let-go* (*t*(12) = 4.39, *p*_FDR_ = 0.004, *d*_*z*_ = 1.22) and accounts for most of the observed difference. In contrast, the *δ, θ*, and *α* bands exhibit small but significant decreases (*p*_FDR_ *≤*0.043, *d*_*z*_ between *™* 0.96 and*™* 0.66), which are, in absolute CSER terms, more than an order of magnitude smaller than the *γ*-band increase. In the *β* band, no significant difference is found. This pattern is also reflected in the aggregated power spectra, where the *let-go* condition generally exhibits increased high-frequency power (≳30 Hz) in most pieces and musicians (see Supplementary Material S1). These findings are also consistent with previous results obtained with different complexity metrics (Dolan et al. 2018).

### B. Higher-order interdependence

Although signal complexity provides valuable information about the dynamical richness of neural activity, it does not explicitly characterise how information is shared and distributed across the interacting components. To study this, we employed two measures that generalise Shannon’s mutual information to multiple variables, namely the S- and O-information (Rosas et al. 2019). S-information captures the overall amount of statistical structure in a set of variables – i.e. how strongly their activity is jointly constrained – providing a natural measure of multivariate interdependence. O-information, in turn, characterises the nature of these dependencies, distinguishing whether they are largely captured by lower-order relationships between the variables (positive O-information), or whether they are only accessible when all variables are considered jointly (negative O-information), i.e. when higher-order dependencies dominate (Rosas et al. 2019).

We applied these estimators to the neural data by computing S- and O-information on triplets of EEG channels. Specifically, for each piece, we considered pairs of musicians and evaluated all possible combinations of channel triplets across the available electrodes. Each triplet is la-belled as either *intra* or *inter*, depending on whether all three channels belong to the same musician or are distributed across both musicians, respectively. For the trio datasets, we repeated this procedure for each of the three pairs of musicians, pooling the resulting triplets.

Most notably, S-information computed on inter-brain triplets is significantly higher in *let-go* than in *strict* for Pieces 1, 2, 4, and 5 (Fig. 3a; *d*_*z*_ between 0.27 and 1.29, e.g. Piece 2: *t*(37) = 4.04, *p*_FDR_ = 0.001, *d*_*z*_ = 0.65), revealing that the activity of the musicians’ brains becomes more strongly interdependent during improvisational performance. This increase is accompanied by a similar rise in intra-brain S-information (significant for Pieces 1, 2, 4, and 5; *d*_*z*_ between 0.21 and 0.95), suggesting that the enhanced coupling between performers goes together with more cohesive activity within each brain. Since the interdependence of neural activity is a central ingredient of theories linking brain integration to conscious experience (Luppi et al. 2019; Tononi 2004; Tononi et al. 1994), its extension across brains provides a natural signature of a jointly structured, shared state. In contrast, O-information shows no consistent *let-go*–*strict* difference, the only significant change being a small decrease of intra-brain O-information in Piece 3 (*t*(59) =*™* 2.49, *p*_FDR_ = 0.035, *d*_*z*_ = *™* 0.32), indicating that the balance between lower- and higher-order dependencies is largely preserved across performance modes. We also note that S-information is consistently higher intra-brain than inter-brain across all pieces and performance modes (Fig. 3b; all *p*_FDR_ *<* 10^*™*3^, *d*_*z*_ between 1.28 and 3.32), as expected given the stronger coupling between electrodes of the same brain.

**FIG 3:**
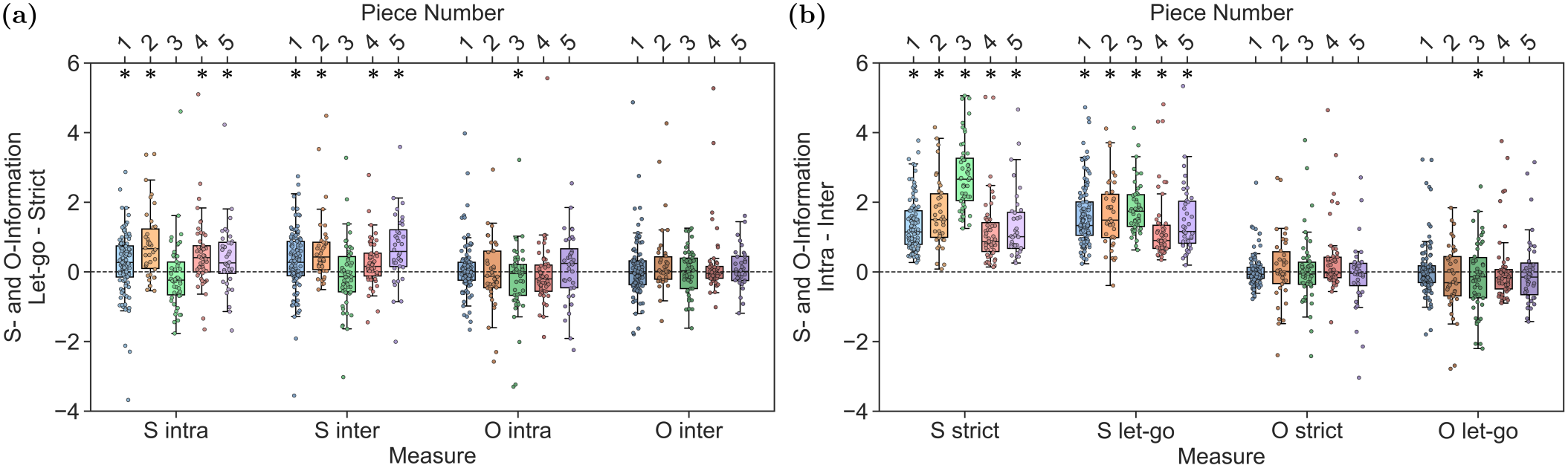
Improvisational performances display stronger interdependence within and between brains. S- and O-information values calculated on all pieces (for the trios, pooling triplets across all pairs of musicians). For each musician, the observed differences are normalised by their standard deviation across electrodes so that each distribution has unit variance. (a) *Let-go*–*Strict* difference, and (b) Intra–Inter difference. *Intra* refers to triplets of channels belonging to the same brain, whereas *Inter* refers to triplets spanning two brains. Stars mark significant differences (one-sample t-test on the paired differences, FDR-corrected *p <* 0.05; see Sec. IV G and Tab. S2).

### C. Directed information-flow networks

We next analysed directed networks derived from Partial Mutual Information from Mixed Embedding (PMIME) (Kugiumtzis 2013), which provides a model-free technique for constructing directed information-flow networks. By applying PMIME to epochs of 1,000 samples across the brain activity of all musicians in each experiment, we obtained a time-resolved sequence of graphs that capture evolving interactions between brain regions, enabling us to study not only within-but also inter-brain interaction dynamics. While information-theoretic approaches provide important insights into the statistical structure of a system, network science offers complementary tools to characterise robustness, community structure, and emergent properties of complex systems. We thus analysed these networks through three lenses: their mesoscale community structure (cluster entropy), their global topology (network measures), and the joint contribution of the musicians’ brains to communication across the network (Partial Network Decomposition, Sec. II D).

#### Cluster entropy

We first examined the community structure of the networks by applying Infomap, a standard flow-based method that partitions each network into groups of nodes (clusters) with stronger internal connectivity (Rosvall et al. 2009). By tracking the evolution of these communities, we obtain a description of how functional brain organisation emerges and reorganises during performance, both within and across musicians. Importantly, since each network comprises the electrodes of all musicians, communities are not restricted to a single brain, and can therefore capture the functional integration of the joint, inter-brain system. We characterised this organisation using cluster entropy, a metric that captures the diversity of community sizes at each time point. Specifically, high cluster entropy suggests a fragmented structure with many small communities, whereas low values correspond to a more coherent organisation dominated by a few large clusters (Fig. 4c).

**FIG 4:**
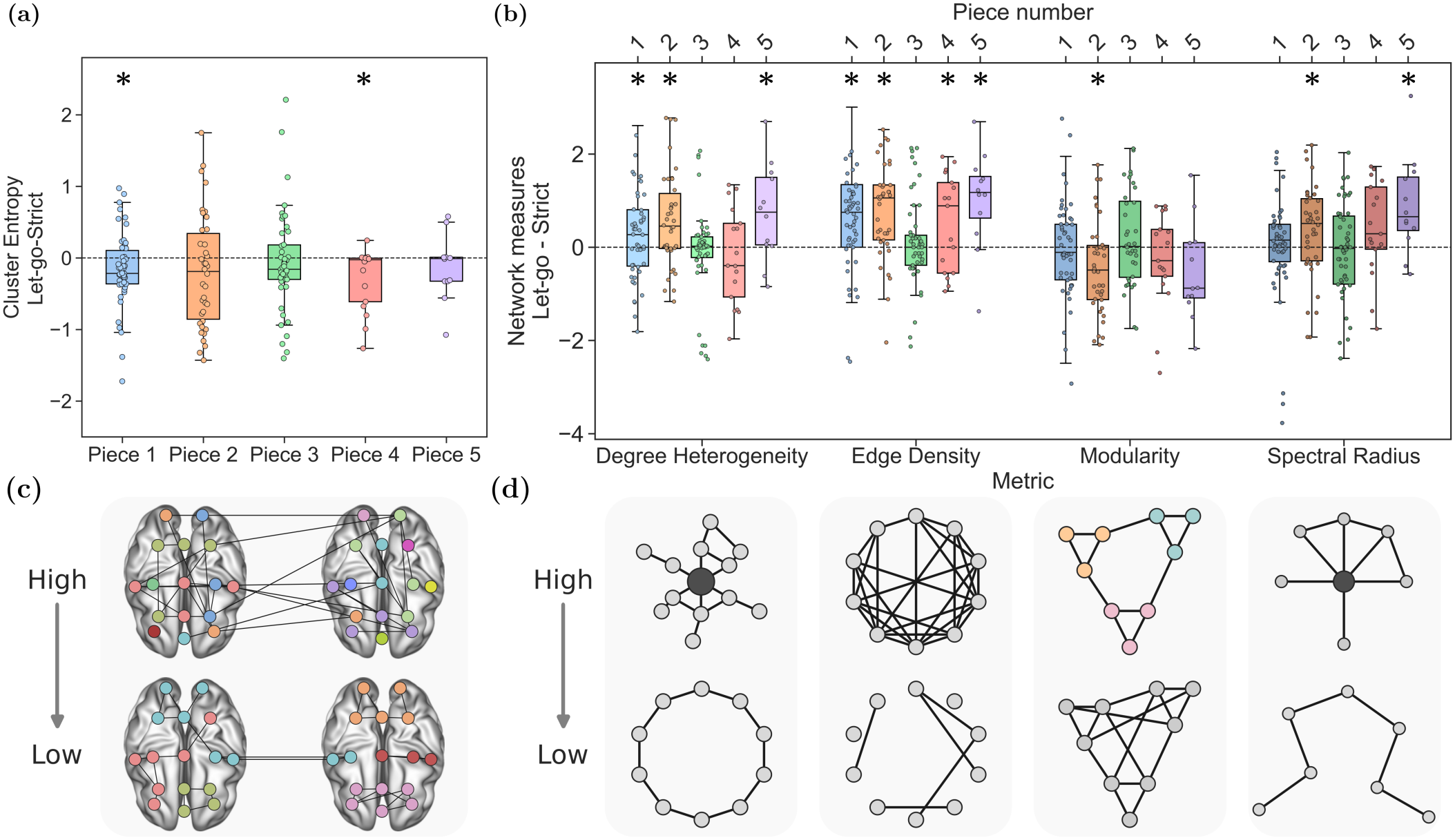
Improvisational performances are characterised by denser and more integrated information-flow networks. (a) *Let-go*–*Strict* differences of cluster entropies calculated on the PMIME networks across all pieces. (b) *Let-go*–*Strict* differences of various network measures across all pieces. Stars mark significant differences (one-sample t-test on the paired differences, FDR-corrected *p <* 0.05; see Sec. IV G and Tab. S3). (c) Illustrative hyperbrain networks (i.e. networks spanning the electrodes of all musicians) with high (top) and low (bottom) cluster entropy, with node colours denoting communities. (d) Illustrative networks with high (top) and low (bottom) values of each measure in (b).

Results show a reduction in cluster entropy during *let-go* (Fig. 4a), which is significant in Pieces 1 and 4 (*t*(110) = *™*3.82, *p*_FDR_ *<* 10^*™*3^, *d*_*z*_ = *™*0.36, and *t*(16) = *™*2.64, *p*_FDR_ = 0.022, *d*_*z*_ = *™*0.64, respectively), suggesting that this performance mode is associated with more integrated network organisation, in agreement with the fewer and larger clusters observed in *let-go* networks (Supplementary Material S3 B). Even when differences do not reach statistical significance, their consistency across heterogeneous datasets points to a general trend, whereby *let-go* is characterised by a more integrated community structure relative to the *strict* condition.

#### Network topology

We further complement these findings through an analysis of standard network properties, focusing on metrics that capture both intercon-nectedness and community structure, including degree heterogeneity, edge density, modularity, and spectral radius (Fig. 4d). A brief description of these measures is provided in Supplementary Material S3, together with the results for additional metrics (Fig. S5). For each experiment, we considered the PMIME networks above and computed these quantities for the *strict* and *let-go* versions of each piece (Fig. 4b).

Overall, the *let-go* mode exhibits higher degree heterogeneity and edge density (significant for Pieces 1, 2, and 5, and for Pieces 1, 2, 4, and 5, respectively; e.g. edge density in Piece 1: *t*(110) = 6.73, *p*_FDR_ *<* 10^*™*8^, *d*_*z*_ = 0.64), indicating more densely connected networks compared to the *strict* condition. This observation is supported by increased weighted clustering, transitivity, and square clustering values, although these reach significance only in a subset of pieces (Supplementary Material S3, Fig. S5, and Tab. S4: weighted clustering in Piece 1, transitivity and square clustering in Piece 2). Additionally, the larger spectral radius observed in *let-go* networks (significant for Pieces 2 and 5) suggests greater resilience and more efficient information diffusion. In contrast, modularity is lower during *let-go* performances (significantly so for Piece 2, *t*(37) = *™*2.58, *p*_FDR_ = 0.020, *d*_*z*_ = *™*0.42), indicating less demarcated community structures. While lower modularity and lower cluster entropy are not equivalent, together with the fewer and larger clusters observed in *let-go* (Supplementary Material S3 B, Fig. S6), these results point towards a less segregated and more integrated network organisation during improvisation.

The analysis of the Flute–Piano–Singer trio dataset (Piece 3) did not yield any significant differences across these measures. This is likely due to the sparsity of networks generated by the PMIME procedure, which in turn results from the narrow frequency range of these EEG recordings, band-limited to 40 Hz at acquisition (Tab. I). We hypothesise that this limited resolution in higher *γ* frequencies reduces the ability to discriminate between *let-go* and *strict* modes, in agreement with the complexity analyses discussed in Sec. II A.

### D. Synergy between brain networks

We conclude our analysis with the application of Partial Network Decomposition (PND) (Luppi et al. 2024a). PND is a framework for quantifying the structural organisation of multiplex networks. Specifically, given two network layers defined on the same set of nodes, PND estimates the synergistic, unique, and redundant contributions of each layer by examining shortest paths between node pairs. A path is classified as *synergistic* if it requires a combination of edges from both layers, *unique* if it relies exclusively on the edges of one layer, and *redundant* if both layers independently provide paths of equal length. These contribution modes are the so-called *PND atoms* (Fig. 5a). Note that this network-level notion of synergy is distinct from the statistical one underlying O-information, as it refers to paths through the network rather than to dependencies among variables.

**FIG 5:**
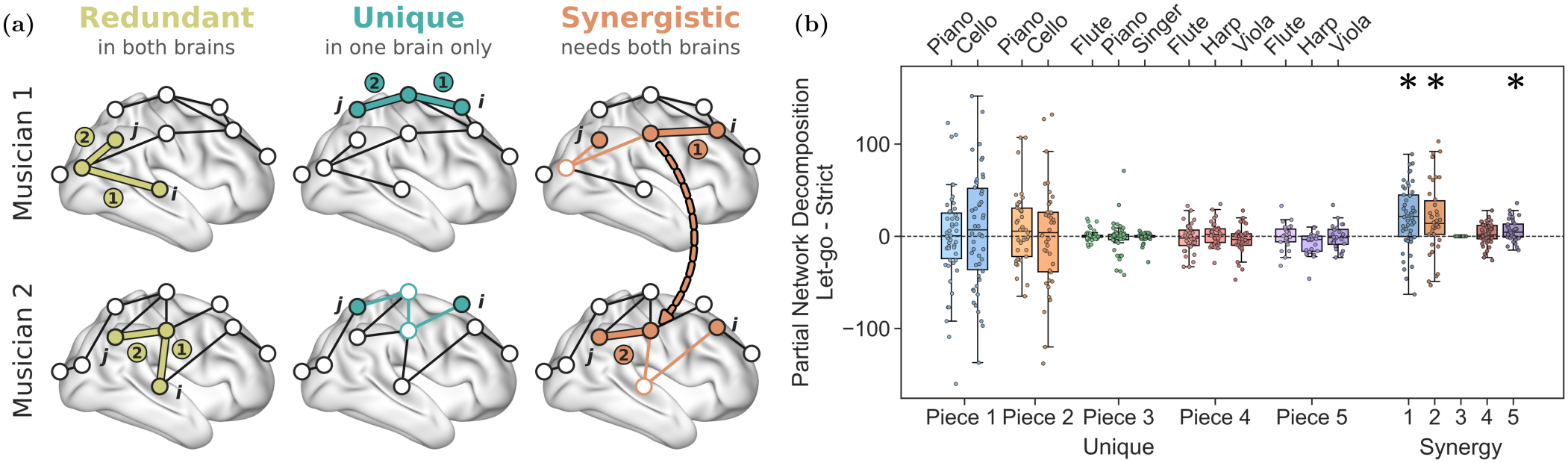
The brain networks of the musicians are more synergistic during improvisational experiences. (a) Schematic of the Partial Network Decomposition approach. The brain networks of two musicians form the two layers of a duplex network defined on the same set of electrodes, so that each layer contains only the intra-brain connections of one musician. Given two nodes *i* and *j*, their shortest path (numbered edges) is *redundant* if each brain independently provides a path of the same length, *unique* if it is found in only one brain (the other offering only a longer route), and *synergistic* if it requires edges from both brains, being shorter than any path available within either brain alone. (b) *Let-go*–*Strict* difference in raw PND atom counts. For the trios, PND is computed on all pairs of musicians, and the results are pooled. Stars mark significant differences (one-sample t-test on the paired differences, FDR-corrected *p <* 0.05; see Sec. IV G and Tab. S5).

Since PND is defined for duplex networks, we applied it to the PMIME networks of each pair of musicians (pooling the results across pairs for the trios), taking the brain networks of the two musicians as the two layers, with nodes matched by electrode position, and comparing the PND atoms between the *let-go* and *strict* conditions. Notably, we observed significantly more synergistic paths in *let-go* than in *strict* for both pieces of the duo and for Piece 5 (Fig. 5b; Piece 1: *t*(110) = 6.60, *p*_FDR_ *<* 10^*™*7^, *d*_*z*_ = 0.63; Piece 2: *t*(37) = 3.14, *p*_FDR_ = 0.015, *d*_*z*_ = 0.51; Piece 5: *t*(11) = 3.17, *p*_FDR_ = 0.027, *d*_*z*_ = 0.91), suggesting that the brain networks of the two musicians become more complementary during *let-go*, as combining their connections provides shorter routes between electrodes than either brain offers alone. Since each layer contains only the intra-brain connections of one musician, these synergistic paths reflect the complementarity of the two brains’ network topologies. This effect is not simply a consequence of higher numbers of connections in the *let-go* networks, as normalising the PND atom counts by the total number of paths preserves the observed patterns (Supplementary Material S3 C, Fig. S7). In Piece 3, very few synergistic paths were found in either condition, consistent with the sparsity of the corresponding networks discussed above. Unique paths show no significant difference, and redundant paths are not reported, as the number obtained was insufficient for meaningful statistical analysis (*<* 10 points).

## III. DISCUSSION

In this work, we investigated the neural correlates of shared experience during musical improvisation using a combination of information-theoretic and network-based approaches. Across most analyses, we observed consistent differences between *let-go* and *strict* performance modes, suggesting distinct underlying modes of functional neural organisation.

### A. Enhanced complexity and cognitive engagement during improvisation

Our analyses revealed a robust increase in signal complexity during the *let-go* condition. Since higher complexity accompanies richer and more vivid conscious states (Blackburne et al. 2025; Casali et al. 2013; Sarasso et al. 2015; Schartner et al. 2017), this increase suggests that each musician’s brain enters a richer and more differentiated state during *let-go*, consistent with the characterisation of the improvisational state of mind as a high-entropy state (Dolan et al. 2018). Moreover, our findings showed that the strongest contribution arises from the *γ* frequency band. The *γ* range has been widely associated with higher cognitive functions, including attentional selection, working memory, and perceptual binding (Fries 2009; Jensen et al. 2007; Tallon-Baudry and Bertrand 1999), as well as being among the first proposed neural correlates of consciousness (Crick and Koch 1990). In this context, the observed increase in *γ*-driven complexity during improvisation suggests a shift towards a more cognitively demanding and integrative brain state, consistent with heightened awareness and flexible information processing relative to the more constrained *strict* condition.

This interpretation is further supported by previous studies linking high-frequency neural activity to complex behavioural and social coordination. In particular, *γ*-band synchronisation has been associated with enhanced performance and richer interpersonal coupling in joint-action and hyperscanning paradigms, whereas lower-frequency dynamics tend to dominate in more con- strained or purely sensorimotor coordination tasks (Barraza et al. 2020; Czeszumski et al. 2020; van Wijk et al. 2012; Wikström et al. 2022). Together, these findings suggest that fast neural dynamics play a central role in supporting flexible, context-sensitive coordination both within and across individuals.

### B. Increased inter-brain coupling and integration in improvisation

S-information was higher in four of the five pieces during *let-go* performances for both intra- and inter-brain interactions. This further indicates that neural activity becomes more structured and interdependent during improvisation, reflecting increased functional integration across distributed processes. The increase in inter-brain S-information is particularly relevant for shared experience, as it reflects a statistical structure that cannot be attributed to either musician alone, but only to their joint activity.

At the hyperbrain network level, multiple complementary measures – including cluster entropy, weighted clustering, transitivity, edge density, and spectral radius – converged on a consistent picture of increased connectivity and more integrated organisation during *let-go* performances. Collectively, these results suggest stronger inter-regional cooperation and more efficient information flow across brain networks.

In particular, reduced cluster entropy and enhanced connectivity metrics suggest a shift towards more coordinated and less fragmented network organisation, consistent with a state of increased functional integration. Since communities in these networks are not restricted to a single brain, this may reflect an increased collective integration across the performers, and not only within each of them. This pattern supports the interpretation that *let-go* performance is characterised by a more globally coupled neural architecture, in which distributed brain regions operate in a more coordinated and interdependent manner. PND further corroborates this view, revealing a higher prevalence of synergistic pathways during *let-go* performances, indicative of more complementary network topologies in the musicians’ brains, which, when combined, provide shorter communication paths than either brain alone. Taken together, these findings provide converging evidence that musical improvisation is associated with stronger neural coordination, more integrated network structure, and increased interdependence both within and across brains. In other words, they support the view that, in *let-go* modes, artistic creation becomes a holistic cognitive experience, characterised by enhanced integration and synergistic processes, and consistent with a measurable neural counterpart of shared experience.

Moreover, these results are in line with previous literature suggesting that musical improvisation relies on dynamic coordination, predictive processing, and flexible integration across distributed brain systems (Daikoku 2024; Ramírez-Moreno et al. 2023; Vergara et al. 2021). Importantly, our work extends these findings by providing a unified quantitative framework that spans information-theoretic, spectral, and network-level descriptions, suggesting that improvisational states are not only behaviourally observable but also detectable in multiscale neural signatures.

### C. Shared experience and signatures of conscious states

Throughout this work, we have adopted a graded view of consciousness, in which complexity, integration, and synergy serve as markers of the richness of a conscious state. This approach is supported by evidence that the same markers discriminate not only between conscious and unconscious states, but also between different conditions within wakefulness. For instance, neural signal diversity distinguishes between conditions of external stimulation, with changes that are associated with subjective ratings of the experience (Liardi et al. 2026; Mediano et al. 2024), and it fluctuates with cognitive engagement even during ordinary wakefulness (Mediano et al. 2021). Altogether, our results show that these signatures are enhanced during *let-go* performances in most pieces. Within each musician’s brain, the increase in signal complexity points to richer and more differentiated activity, while the increase in intra-brain S-information indicates more integrated activity. The co-occurrence of these two effects is notable, as conscious states have long been characterised as being at once highly differentiated and highly integrated (Tononi and Edelman 1998; Tononi et al. 1994), and complexity has emerged as a common denominator of consciousness across very different regimes of brain dynamics (Frohlich et al. 2022; Sarasso et al. 2021). Across brains, the increase in inter-brain S-information, the more integrated hyperbrain communities, and the larger number of synergistic paths obtained by combining the two brains’ networks indicate that the joint system formed by the musicians displays the same signatures of integration and synergy that, within single brains, accompany conscious states (Luppi et al. 2022, 2024b; Tagliazucchi et al. 2016). Within this framework, these results suggest that *let-go* performances correspond to richer conscious states, not only within each musician but also in the joint system they form.

These findings also offer a possible neural counter-part to the phenomenology of shared experience, which is characterised by a sense of unity with others (Chung et al. 2024) and a blurring of the boundary between self and other (Tarr et al. 2014). During *let-go*, the musicians’ brains behave less like separate systems exchanging information and more like parts of a single integrated system, in which fewer and larger communities can span different brains and the network of each brain increasingly complements that of the other. Notably, reduced within-network integrity and increased between-network connectivity have been correlated with ego dissolution under psychedelics within individual brains (Lebedev et al. 2015; Nour and Carhart-Harris 2017; Tagliazucchi et al. 2016), suggesting that a loosening of functional boundaries may be a common feature of experiences in which the sense of self extends beyond the individual.

Shared experiences are closely tied to *theory of mind* and our ability to infer others’ mental states, enabling groups to coordinate and collaborate more effectively in joint problem-solving (Engel et al. 2014). Such interpersonal cognition underpins *collective intelligence*: a group’s capacity to collaboratively solve problems and create ideas beyond individual abilities (Leimeister 2010; Woolley et al. 2015), resulting in novel group dynamics such as risk-taking (Wang et al. 2019), action coupling (Woolley et al. 2010) and collective adaptation (Galesic et al. 2023). Our analysis can thus provide a research paradigm for studying the phenomenological aspects underlying collective intelligence.

### D. Limitations and future steps

Musical performance, although a useful naturalistic paradigm, is also radically heterogeneous – spanning many instruments, genres, and ensemble configurations. Thus, our limited set of performers and experimental conditions may not generalise to other musicians or performance environments. Additionally, the resolution of the EEG data, particularly at high *γ* frequencies, constrained the detection of subtle network effects, especially in the trio datasets, which were recorded with lower sampling rates or narrower frequency ranges, and, for the Flute– Harp–Viola trio, fewer channels (Tab. I). Moreover, each piece was performed only once in each mode, and our statistical comparisons are carried out across epochs within these performances. While the consistency of the results across ensembles and pieces supports their robustness, larger samples of performers and repeated performances would be needed to generalise them at the population level. A possible additional limitation is that intra-brain dependencies may be inflated by volume conduction, as the same neural source can be picked up by several electrodes. However, since this effect depends on head anatomy and electrode montage, which are unchanged across conditions, we expect it to largely cancel out when contrasting *let-go* and *strict* performances. This cannot affect inter-brain interactions, as channels on different scalps share no conducted source. Finally, *let-go* performances were accompanied by more pronounced body movements and facial expressions, which can introduce muscle artefacts predominantly in the high-frequency range. Although we adopted a dedicated pipeline to remove muscle activity (Sec. IV F), differences in movement between conditions represent an important consideration when interpreting high-frequency activity.

Future studies could strengthen and extend these findings by including a more diverse sample of performers, employing a denser EEG headset, or analysing longer and more numerous musical pieces. Such efforts would improve statistical power, allow for more detailed characterisation of higher-order interactions, and test the generality of the markers identified here. Moreover, combining these approaches with behavioural measures and subjective reports (e.g. post-performance interviews, or continuous self-reports such as real-time tapping of perceived connection) could further clarify how neural dynamics relate to musical creativity and collaborative experience.

### E. Conclusion

Overall, our study suggests that the shared experience of group improvisation leaves measurable signatures in the musicians’ brains and, crucially, in the interactions between them. In this work, we detected them through a variety of mathematical tools, from information-theoretic measures to network-based estimators. Across these frameworks, we found markers consistent with shared experience in most pieces, as captured by higher intra- and inter-brain S-information, increased network connectivity and clustering, and a more synergistic combination of the musicians’ brain networks. These findings indicate that *let-go* performances are associated with greater coordination across brain regions, highlighting the emergence of an integrated, holistic neural state during creative activity.

Importantly, beyond revealing these neural signatures, this work also provides a set of tools and methodologies that can be applied to hyperscanning studies. The combination of complexity measures, information-theoretic analyses, and network-based metrics offers a general framework to investigate inter-brain coordination, synergy, and network-level interactions across diverse experimental settings.

In sum, this work yields both empirical evidence and computational tools to characterise improvisation as not only an individual cognitive state, but also a collective one, marked by enhanced interdependence, cooperation, and synergy between neural systems. These findings advance our understanding of the neural foundations of shared experience, suggesting that the informational signatures of conscious states may extend beyond individual brains to interacting systems, while providing quantitative markers for investigating the collective dimensions of conscious experience, artistic collaboration, and improvisational expertise.

## IV. METHODS AND DATA

### A. Complexity via State-space Entropy Rate

Complexity via State-space Entropy Rate (CSER) is an information-theoretic measure of signal complexity grounded in the framework of state-space models (Mediano et al. 2023). State-space (SS) models provide a flexible and robust approach for modelling time series data, describing the evolution of an unobserved (latent) dynamical system that generates the observed signals. This framework is widely used in the study of multivariate and stochastic processes, with applications to signal processing (Larimore 1990), control the-ory (Åström 2012; Kalman 1960), computational neuro-science (Paninski et al. 2010; Smith and Brown 2003), and beyond (Durbin et al. 2004; Harvey 1990), as it enables separating the underlying dynamics from observation noise.

A general linear state-space model is defined by two equations:

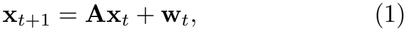

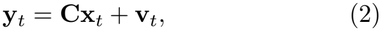

where **x**_*t*_ denotes the latent state vector, **y**_*t*_ the observed data, **A** the state transition matrix, and **C** the obser-vation matrix. The terms **w**_*t*_ and **v**_*t*_ represent process and observation noise, respectively. In this work, we restrict attention to the Gaussian case, where both noise terms are assumed to be zero-mean multivariate Gaussian processes with covariance matrices **Q** and **R**, and cross-covariance **S**.

Under Gaussian assumptions, the state-space model can be equivalently expressed in innovation state-space (ISS) form, where observations are written in terms of a one-step-ahead prediction error (innovation) process. More precisely, by introducing the quantities 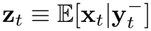 and 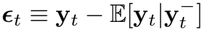, where 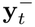 denotes the past of the observed process, the ISS representation is given by

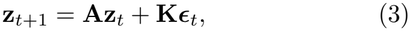

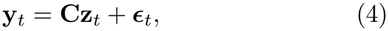

where **z**_*t*_ denotes the optimal state estimate (i.e. the Kalman filter state), ***ϵ***_*t*_ is the innovation process, and **K** is the Kalman gain matrix. The innovation process ***ϵ***_*t*_ is a zero-mean Gaussian white noise with covariance matrix **V**, and it captures the part of the signal that is unpredictable from its past.

With this framework at hand, CSER is then defined as the entropy rate of this innovation process, which for a Gaussian process depends solely on its covariance:

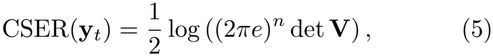

where **V** is the covariance matrix of the innovations and *n* is the dimensionality of the observed process.

CSER has been applied to neural data, particularly in the study of different levels of consciousness, where it provides a robust and interpretable measure of dynamical richness (Liardi et al. 2026; Mediano et al. 2023). In this work, for each piece in all datasets, we segmented the EEG signal into epochs of 1,000 samples and computed CSER for each channel.

### B. S- and O-information

S- and O-information are information-theoretic quantities designed to characterise higher-order statistical dependencies in multivariate systems (Rosas et al. 2019). Unlike pairwise measures such as mutual information, which only capture dependencies between two variables at a time, these quantities capture dependencies involving three or more variables, thereby providing access to collective structure, with O-information in particular revealing whether higher-order dependencies dominate. These metrics have been successfully applied across a wide range of domains, including neural (Faes et al. 2022; Varley et al. 2023) and physical systems (Caprioglio et al. 2025; Orio et al. 2023; Robiglio et al. 2025).

To present S- and O-information, we first introduce the total correlation (TC), also known as multi-information (Watanabe 1960), which is defined as

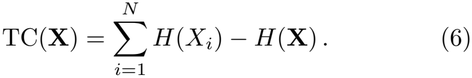

The TC quantifies the total amount of statistical dependence among the variables **X** = (*X*_1_, … , *X*_*N*_), or, in other words, it captures how much the joint distribution deviates from independence. These dependencies are also referred to as *collective constraints*.

A complementary quantity which aims to explore multivariate dependencies from a different angle is the dual total correlation (DTC) (Te Sun 1978), which reads

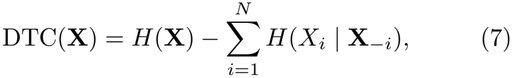

where **X**_*™i*_ denotes all variables except *X*_*i*_. DTC measures the part of the joint entropy that is shared by at least two variables, i.e. the joint entropy minus the uncertainty that is private to each individual variable.

Since TC and DTC provide complementary measures of interdependence, the S- and O-information combine them to disentangle the overall strength of dependencies from their qualitative nature:

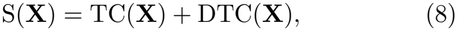

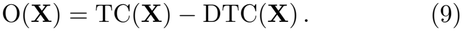

S-information quantifies the overall magnitude of statistical structure in the system, due both to collective constraints and shared randomness. In contrast, the O-information estimates the balance between redundancy and synergy. More specifically, positive values of O-information indicate redundancy-dominated dependencies, whereas negative values indicate predominantly synergistic regimes.

In our analysis, we studied S- and O-information on triplets of channels across musicians’ brains, estimated under Gaussian assumptions on epochs of 1,000 samples. The choice of triplets is motivated by two considerations: first, higher-order concepts such as synergy and redundancy are more readily interpretable in low-dimensional settings; second, restricting the analysis to triplets simplifies the distinction between *intra* and *inter* configurations, avoiding the combinatorial complexity that would arise with larger sets of electrodes.

### C. Partial Mutual Information from Mixed Embedding

Partial Mutual Information from Mixed Embedding (PMIME) is a model-free, information-theoretic method for detecting directed interactions in multivariate time series (Kugiumtzis 2013). It is specifically designed to infer direct causal influences while mitigating the effects of indirect couplings and common drivers, which often confound traditional pairwise measures. This technique has been applied to a variety of real-world datasets, including physiological and neural recordings (Kugiumtzis et al. 2017), financial markets (Fotiadis et al. 2023), and environmental systems (Ren et al. 2025).

PMIME operates by constructing a mixed embedding vector that iteratively selects lagged components from all variables in the system. At each step, the algorithm identifies the candidate component that maximises the conditional mutual information with the target variable, given the already selected components. This forward selection procedure continues until a termination criterion is met, i.e. until the additional information contributed by the selected component becomes negligible relative to the information already explained, resulting in a parsimonious embedding that captures the relevant dynamical dependencies.

Formally, let *Y*_*t*+1_ denote the future state of the target variable, and 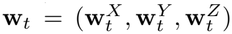 the resulting mixed embedding vector, partitioned into the lagged components selected from the source *X*, from the target *Y* itself, and from all remaining variables *Z*. PMIME estimates the directed influence from a source variable *X* to *Y* as

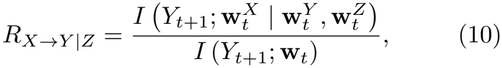

i.e. the fraction of the information about the future of *Y* explained by the mixed embedding that is uniquely contributed by *X*. By construction, *R*_*X→Y* |*Z*_ *∈* [0, 1], and it is exactly zero whenever no component of *X* is selected in the embedding. All (conditional) mutual information terms are estimated with the nearest-neighbour estimator of Kraskov et al. (2004). By conditioning on the mixed embedding, PMIME isolates direct effects and reduces spurious detections. By repeating this procedure for each target variable, one can construct a causal connectivity matrix that describes the statistical inter-dependencies across all components of the system over time.

In this work, we used the MATLAB implementation of PMIME by Kugiumtzis^1^ with its default parameters, applying it separately to each epoch and including all EEG channels of all musicians as variables. Specifically, candidate components comprise the current and the four previous samples of each variable (*L*_max_ = 5), the embedding aims at explaining the target one step ahead (*T* = 1), and all information-theoretic quantities are estimated with *k* = 5 nearest neighbours. The embedding terminates when the ratio *I*(*Y*_*t*+1_; *w*|**w**_*t*_)*/I*(*Y*_*t*+1_; *w*, **w**_*t*_) between the conditional mutual information of the newly selected component *w* and the mutual information of the augmented embedding falls below *A* = 0.03, or when this ratio increases over two consecutive embedding cycles, with a maximum of 20 components. Before the analysis, each time series is rescaled to the unit interval. The resulting matrix of *R*_*X→Y* |*Z*_ values defines a directed network over the electrodes of all musicians, which we use directly as a weighted adjacency matrix.

### D. Cluster entropy

Let 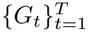 be a set of graphs evolving over time, where *G*_*t*_ = (*V, E*_*t*_) denotes the network at time *t*, with node set *V* and edges *E*_*t*_. At each time point, partitioning *G*_*t*_ via a community detection algorithm provides a set of communities 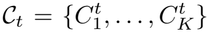. In this work, communities were identified with Infomap (Rosvall et al. 2009), which partitions a network into modules by minimising the description length of a random walk on the network (the so-called map equation). We used the Python implementation of Infomap (Edler et al. 2023, v. 2.9.2), taking the PMIME matrices directly as weighted adja-cency matrices. We searched for a two-level partition, i.e. a flat partition into non-overlapping modules without hierarchical sub-modules, and retained the partition with the shortest description length across 200 independent trials. All other parameters were left at their default values, namely Markov time 1, no regularisation, and a maximum of 10 core loops per optimisation level.

For each community 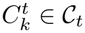, we then define the probability

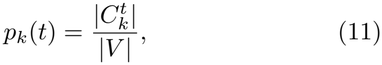

corresponding to the fraction of nodes assigned to that community at time *t*, i.e. the probability that a randomly chosen node at time *t* belongs to community *k*. We then define the *cluster entropy* as the Shannon entropy of this distribution:

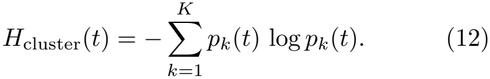

This quantity summarises the mesoscale organisation of the network at each time point. High values of *H*_cluster_(*t*) indicate a more fragmented structure, characterised by many small communities of comparable size and therefore higher uncertainty in the assignment of nodes to clusters. Conversely, low values correspond to more organised networks dominated by a few large communities, reflecting a more cohesive, integrated organisation.

### E. Partial Network Decomposition

Partial Network Decomposition (PND) is a framework that quantifies how different layers or components of a multiplex network contribute to a network-level property of the system (Luppi et al. 2024a). This approach is particularly useful in the analysis of complex systems, where interactions occur across multiple modalities or scales. Examples of these scenarios can be found in neuroscience, where brain networks might combine structural and functional connectivity (Lim et al. 2019) or different modes of information processing (Rajpal et al. 2026), as well as in social sciences, where layers may represent different relations among people (Nakis et al. 2026). PND has been employed in the study of multiplex networks in mammalian brains and public transportation systems (Luppi et al. 2024a).

Leveraging a mapping between graph-theoretic quantities and expectations over node pairs, PND provides a framework to attribute the value of a network-level observable to different layers of a multiplex network. Specifically, a network property (e.g. global efficiency or shortest path length) is expressed as the average of a pairwise function over all node pairs, and the contribution of different layers is assessed by comparing how this quantity changes when edges from different layers are considered alone or jointly. This construction enables a decomposition of the observable into redundant contributions (shared across layers), unique contributions (specific to individual layers), and synergistic contributions (emerging only when layers are combined). In this way, PND extends ideas from partial information decomposition (Williams and Beer 2010) to network analysis, allowing one to quantify how different connectivity layers jointly and nonlinearly shape the system-level property of interest. In our application, the two layers are the intra-brain PMIME networks of a pair of musicians, with nodes identified by electrode position. In this regard, PND quantifies how the network topologies of the two brains combine.

### F. Experimental procedure and data cleaning

For the following analyses, we consider three independent musical experiments: a novel one, presented here for the first time, and two that have been studied previously (Dolan et al. 2018, 2013; Wan et al. 2014). The first dataset comprises a performance by the duo of Professor David Dolan (pianist) and Professor Adrian Brendel (cellist), who played two classical pieces (Pieces 1 and 2). The second consists of a Schubert piece (Piece 3) performed by a professional trio, which included Rosie Bowker (flautist), Thibault Charrin (pianist), and Kate Smith (singer) (Dolan et al. 2018). Finally, the third dataset comprises a musical performance of two pieces (Pieces 4 and 5) played by Trio Anima, a classical chamber music group composed of Matthew Featherstone (flautist), Anneke Hodnett (harpist), and Drew Balch (violist) (Dolan et al. 2013; Wan et al. 2014). All the pieces mentioned were played twice, once in the *strict* mode and once in the *let-go* mode, allowing for consistent comparisons between the two.

In the following sections, we provide a detailed description of the data acquisition and preprocessing for the first dataset, while we present a brief overview of the other two, referring to the original papers for a more exhaustive discussion. Importantly, here we only focus on the EEG data of the performers. For all datasets, after the preprocessing steps described below, the signals of each channel were z-scored before any analysis.

#### 1. Piano–Cello duo

##### Experiment description

On 18 June 2019, the two musicians participated in an experiment taking place at the Blyth Centre for Music and Visual Arts at Imperial College London, in collaboration with the Departments of Medicine, Computing, Mathematics, and the Data Science Institute at Imperial College. The performance took place in a dedicated session in front of a small audience of approximately ten people from diverse backgrounds, with audio and video recordings produced by one video camera placed in front of each musician and by semi-professional audio equipment. Neural data were recorded at a sampling frequency of 250 Hz using two gTec EEG caps with 16 and 64 channels, placed on the cellist’s and pianist’s scalps, respectively. However, for symmetry, only 16 of the 64 channels of the second cap were connected. The electrodes were positioned following the standard 10–20 montage.

The experiment consisted of the performance of two pieces, the first by Beethoven (Piece 1) and the second by Schumann (Piece 2). Both pieces were played twice, in *strict* and *let-go* mode.

##### Data preprocessing

The musicians’ extensive movements during the performance resulted in heavily artefactual signals, especially during the *let-go* interpretations. In fact, a previous study has already shown the presence of fast movements during improvised per-formances (Dolan et al. 2018). We confirmed this by inspecting the video recordings, which showed substantial head movements and facial expressions, especially in the case of the cellist. This led to brain signals with heavy eye and muscle artefacts that can mask the neural activity, posing strong constraints on the cleaning procedure: removing artefactual components while preserving as much neural signal as possible was particularly challenging with only 16 channels, and without electrooculogram (EOG) and electromyogram (EMG) recordings.

We first high-pass filtered the data at 1 Hz and manually removed bad channels and epochs by visual inspection of the data. As notch filters negatively affect information-theoretic studies (Barnett and Seth 2011), the line noise at 50 Hz was removed by a combination of Zapline-plus (de Cheveigné 2020; Klug and Kloosterman 2022) and Cleanline (Mullen 2012) techniques. These aim to remove the 50 Hz and higher harmonic components while leaving the background signal uncorrupted, and have been shown to perform better when employed together (Miyakoshi et al. 2021). As anticipated above, the extent of muscle contamination, together with the small number of channels available, precluded removing it with Independent Component Analysis (ICA) alone, as too many components would have had to be excluded, resulting in a significant loss of neural signal. Hence, we employed Removal of Muscle artefacts from EEG (ReMAE) (Chen et al. 2019a), a toolbox implementing state-of-the-art signal-processing techniques for the removal of fast-oscillating muscle activity. Among the variety of ReMAE’s methods, we chose the EEMD-CCA approach (Chen et al. 2019b), consisting of performing Ensemble Empirical Mode Decomposition (EEMD) (Wu and Huang 2009), a data-driven signal decomposition that partitions the signal into spectrally independent fast and slow oscillations, followed by a Canonical Correlation Analysis (CCA) (De Clercq et al. 2006), a Blind Source Separation (BSS) model that forces the reconstructed sources to be maximally autocorrelated and mutually un-correlated. By combining the two techniques, we used EEMD to generate multiple time series for each input, thereby overcoming the limitation of performing BSS on only 16 channels. We then exploited the fact that muscle activity typically spans a wide range of frequencies, resulting in lower autocorrelations than neural signals, thus enabling us to eliminate components with very low autocorrelations via CCA. EEMD-CCA has been applied to EEG data to remove both eye and muscle artefacts, proving its effectiveness in various recent studies (Chen et al. 2017; Yang et al. 2017).

Since performing a BSS procedure on a large number of widely diverse components could be more detrimental than beneficial, following the suggestions of the original paper (Chen et al. 2019a), after the EEMD, we set a threshold on the autocorrelation, so that only components with lower autocorrelation than the selected thresh-old would be included in the CCA procedure. Then, the CCA sources whose autocorrelation remains below a second (rejection) threshold were discarded. To fix these thresholds, we inspected the autocorrelation distribution of the EEG post-EEMD (Supplementary Material S2, Fig. S4), where we noticed that there are typically two groups of components, with low and high autocorrelation respectively, separated by a clear gap. Components affected by muscle activity are expected to fall in the former group. After performing this investigation on different epochs and pieces, we set both selection and rejection thresholds at 0.5, hence only removing the extremely poorly autocorrelated components. We underline that this conservative approach was adopted to minimise the amount of neural data removed. Moreover, potential remaining artefacts can still be addressed during the ICA procedure.

After removing most muscle artefacts, we re-referenced the signal to the average and performed the ICA on the resulting data, manually eliminating residual artefactual components. Lastly, we spherically interpolated the bad channels excluded at the beginning. Although many different re-referencing techniques have been proposed (Bigdely-Shamlo et al. 2015; Yao 2001), we left the signal average-referenced as it is often more suitable for whole-brain studies (Lei and Liao 2017).

Repeating this procedure for both performers and in all EEG recordings, we performed cleaning steps and ICA-component removal as similarly as possible across pieces. However, since for each piece bad epochs for the cellist and the pianist would not coincide in general, we removed the union of the bad segments of both musicians from both recordings to maintain synchronisation between the two brains.

#### 2. Flute–Piano–Singer trio

The trio played a section of Franz Schubert’s *Der Hirt auf dem Felsen* (“The Shepherd on the Rock”) twice, following the paradigms of *strict* and *let-go* modes. Each artist had an EEG cap with 19 electrodes (Fp1, Fp2, F3, F4, F7, F8, C3, C4, T7, T8, P3, P4, P7, P8, O1, O2, Fz, Cz, Pz), placed according to the standard 10–20 positioning system and sampled at 250 Hz. A band-pass filter was applied between 2 and 40 Hz, and bad channels and epochs were visually removed. We refer to Dolan et al. (2018) for a more thorough description.

#### 3. Flute–Harp–Viola trio

Musicians played both purely improvised pieces and composed music. Specifically, the order of the performance was the following:

- Ibert (*strict* and *let-go*)
- Telemann (*let-go* and *strict*)
- Improvisation (*let-go* and *strict*)
- Ravel (*strict* and *let-go*)
- Improvisation (*let-go* and *strict*)

EEG data were recorded from 10 electrodes (P4, O2, T8, C4, F4, F3, C3, T7, P3, O1) at a sampling frequency of 100 Hz. Of the five pieces played, we focus on the last two, as they are the only ones long enough after cleaning to provide meaningful statistics. Data preprocessing consisted of high-pass filtering at 1 Hz and visual removal of bad epochs affected by artefacts. Further information can be found in Dolan et al. (2013) and Wan et al. (2014).

### G. Statistical analysis

All comparisons between the *let-go* and *strict* performances of a piece were paired, as the two versions are renditions of the same musical material by the same musicians. For each measure, we computed the paired differences (*let-go™ strict*) and tested whether their mean differed from zero with a one-sample t-test.

The pairing unit depended on the measure. For broad-band CSER, the samples were EEG channels, pairing the same electrode across the two conditions, and each musician and piece was tested separately; in addition, we averaged broadband CSER over channels and tested it across all 13 musician–piece combinations, as a single test of the overall effect. For the frequency-resolved CSER, band-limited values were averaged over channels, yielding one value per musician and piece in each condition, and each frequency band was then tested across all 13 musician–piece combinations. For the epoch-wise measures (S- and O-information, network measures, cluster entropy, and PND atoms), the samples were epochs, paired by their position within the piece. Epoch pairs in which a measure was undefined in either condition (e.g. clustering coefficients and modularity of networks with fewer than 10 edges) were excluded. For the trios, pair-wise measures (S- and O-information and PND atoms) were averaged across the pairs of musicians within each epoch, so that each epoch contributed a single observation. Finally, the comparison between intra- and inter-brain S- and O-information was paired by epoch within each performance mode.

Tests were one-sided whenever we had an *a priori* hypothesis on the direction of the effect, namely higher values in *let-go* for broadband CSER, S-information, PND synergy, edge density, degree heterogeneity, spectral radius, weighted clustering, transitivity, square clustering, and maximum cluster size; lower values in *let-go* for modularity, cluster entropy, and number of clusters; and higher intrathan inter-brain S-information in both performance modes. All the remaining tests (band-limited CSER, O-information, unique PND atoms, and graph heterogeneity) were two-sided. We controlled for multiple comparisons with the Benjamini–Hochberg false discovery rate (FDR) procedure (Benjamini and Hochberg 1995), applied within each figure panel, i.e. across all pieces, musicians, and measures shown together, and considered results significant for FDR-corrected *p <* 0.05. As a measure of effect size, we report Cohen’s *d*_*z*_, i.e. the mean of the paired differences divided by their standard deviation, which is the standardised effect evaluated by the paired test (Lakens 2013).

Throughout the paper, we report the *t*-statistic with its degrees of freedom, the FDR-corrected p-value (*p*_FDR_), and *d*_*z*_. The complete results are reported in Supplementary Material S4, together with a non-parametric validation based on sign-flip permutation tests, in which the outcome differs from the t-test in only three borderline cases.

### H. Ethics and consent

The experiments on the two trios were conducted as part of previous studies. The experimental protocol for the Flute–Harp–Viola trio received ethical approval from the Guildhall School of Music and Drama (Wan et al. 2014). For the Flute–Piano–Singer trio, informed consent was obtained through a letter of invitation to all participants outlining the experimental procedure and asking them to confirm their acceptance, and performers gave explicit written permission for their identity to be revealed (Dolan et al. 2018). The experiments reported in our manuscript were part of the protocol approved by the Ethics Committee of Guildhall School of Music and Drama. A separate ethics approval for the research reported in our manuscript was not required as per Imperial College Research Ethics Committee’s guidelines as well as national regulations. For the Piano–Cello duo, both musicians were informed about the experimental procedure and provided written informed consent prior to participation, including explicit permission for their identity to be revealed.

## ACKNOWLEDGEMENTS

We are deeply grateful to Professor David Dolan (pianist) and Professor Adrian Brendel (cellist) for their generosity with their time during the performances and for their deep insights during the discussion sessions. Furthermore, we would like to thank all the musicians involved for their enthusiasm, patience, and time. Without the commitment of Matthew Featherstone (flautist), Anneke Hodnett (harpist), Drew Balch (violist), Rosie Bowker (flautist), Thibault Charrin (pianist), and Kate Smith (singer), our research would not have been possible.

## DATA AVAILABILITY

The EEG data of the Piano–Cello duo collected in this study are available from the authors upon request. The trio datasets were collected in previous studies (Dolan et al. 2018, 2013; Wan et al. 2014).

## AUTHOR CONTRIBUTIONS

Conceptualisation, A.L., M.I.S., H.R., and H.J.J.; data curation, A.L., M.I.S., and I.-F.B.; formal analysis, A.L. and M.I.S.; funding acquisition, P.A.M.M., F.E.R., and H.J.J.; investigation, A.L.; project administration, A.L. and H.J.J.; methodology, A.L. and H.J.J.; software, A.L.; supervision, P.A.M.M. and H.J.J.; writing – original draft, A.L. and M.I.S.; writing – review & editing, A.L., M.I.S., H.R., I.-F.B., F.E.R., P.A.M.M., and H.J.J.

## Footnotes

1 https://github.com/dkugiu/Matlab

