## Supplementary Information for "Network Correlates of Shared Experience in Musical Improvisation"

### S1. POWER SPECTRA

Here, we provide the power spectral densities (PSDs) for all pieces and musicians across all datasets.

For each musician, EEG channel, and performance mode, we estimated the PSD with Welch's method, using the whole continuous preprocessed recording of each performance. Unlike in the other analyses, the data were neither split into epochs nor normalised, so that differences in absolute power between conditions are preserved. Each recording was divided into segments of 2048 samples with 50% overlap; each segment was demeaned and tapered with a Hann window, and the resulting periodograms were averaged to obtain a one-sided spectral density, expressed in $\mu V^2/Hz$. A segment lasts 8.2 s at 250 Hz (Pieces 1–3) and 20.5 s at 100 Hz (Pieces 4–5), giving a frequency resolution of approximately 0.12 Hz and 0.05 Hz, respectively.

For each musician and condition, the channel spectra are summarised by their mean across channels (solid lines) and by their minimum and maximum across channels at each frequency (shaded regions), comparing the *let-go* and *strict* modes (Figs. S1–S3). Spectra are shown on a logarithmic scale, up to 100 Hz for Pieces 1–2, 40 Hz for Piece 3 (the upper limit of the acquisition band, see Tab. I of the main text), and 50 Hz, i.e. the Nyquist frequency, for Pieces 4–5.

*

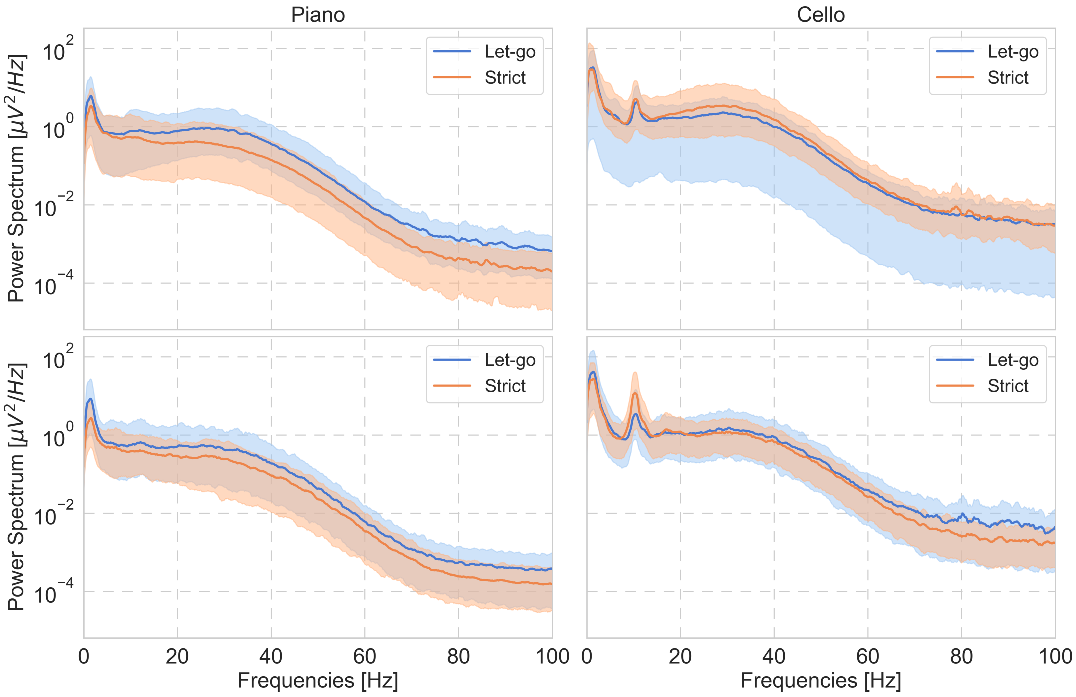


**FIG. S1:** PSDs for piano (left) and cello (right) during Pieces 1 and 2 (first and second rows, respectively). The solid lines indicate the averages across all channels, while the shaded regions show the range (minimum–maximum) across channels.

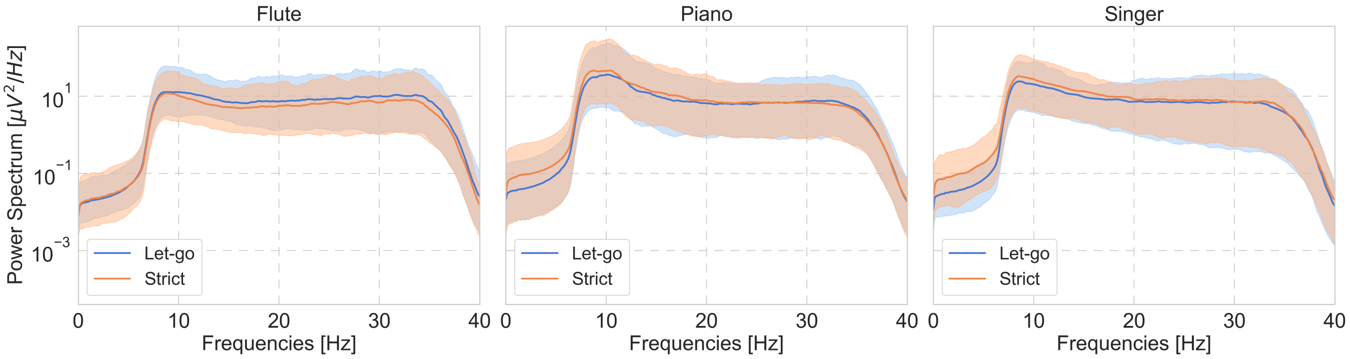


**FIG. S2:** PSDs for flute, piano, and singer (from left to right) of the Flute–Piano–Singer trio (Piece 3). The solid lines indicate the averages across all channels, while the shaded regions show the range (minimum–maximum) across channels.

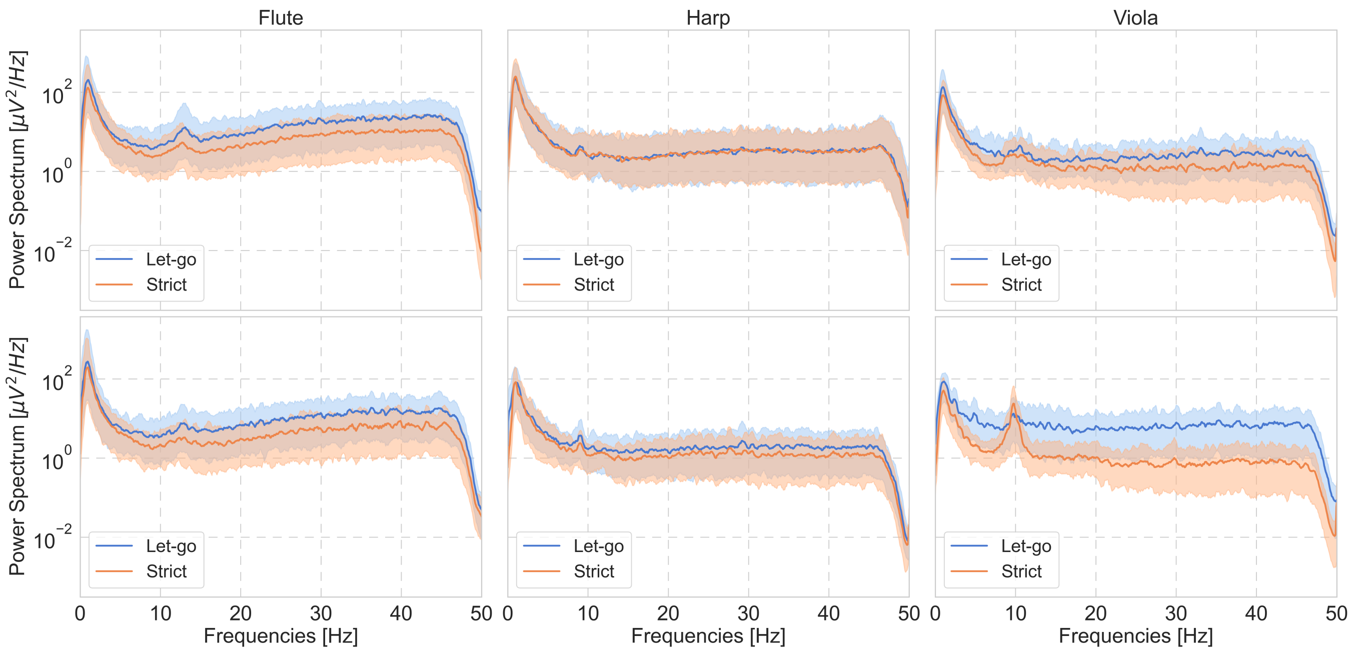


**FIG. S3:** PSDs for flute, harp, and viola (from left to right) of the Flute–Harp–Viola trio during Pieces 4 (first row) and 5 (second row). The solid lines indicate the averages across all channels, while the shaded regions show the range (minimum–maximum) across channels.

### S2. EEMD SIGNAL AUTOCORRELATION

As described in Sec. IV F of the main text, muscle artefacts in the recordings of the Piano–Cello duo were removed with the EEMD-CCA approach (Chen et al. 2019a,b). Ensemble Empirical Mode Decomposition (EEMD) (Wu and Huang 2009) first decomposes each EEG channel into a set of components with different characteristic frequencies, and Canonical Correlation Analysis (CCA) (De Clercq et al. 2006) is then applied to these components to obtain sources that are maximally autocorrelated and mutually uncorrelated. Since muscle activity is broadband, it yields components with low autocorrelation, whereas neural activity is comparatively slow and strongly autocorrelated. The procedure requires two thresholds on the autocorrelation: a selection threshold, so that only the EEMD components below it enter the CCA, and a rejection threshold, below which the resulting CCA sources are discarded.

To set these thresholds, we inspected the distribution of the autocorrelation of the EEMD components. Fig. S4 shows a representative example from our preprocessing pipeline. Most components have autocorrelation above approximately 0.75, clustered around 0.8 and 1, while two smaller groups have much lower values (around −0.3 and 0.3). These groups are separated by a clear gap, with no components between approximately 0.45 and 0.75. We observed a similar separation across epochs and pieces, and therefore set both thresholds at 0.5, within the gap. Only the poorly autocorrelated components, which are the ones most likely affected by muscle activity, were thus removed. This conservative choice minimises the loss of neural signal, and any residual artefacts were addressed in the subsequent ICA step (see Sec. IV F of the main text).

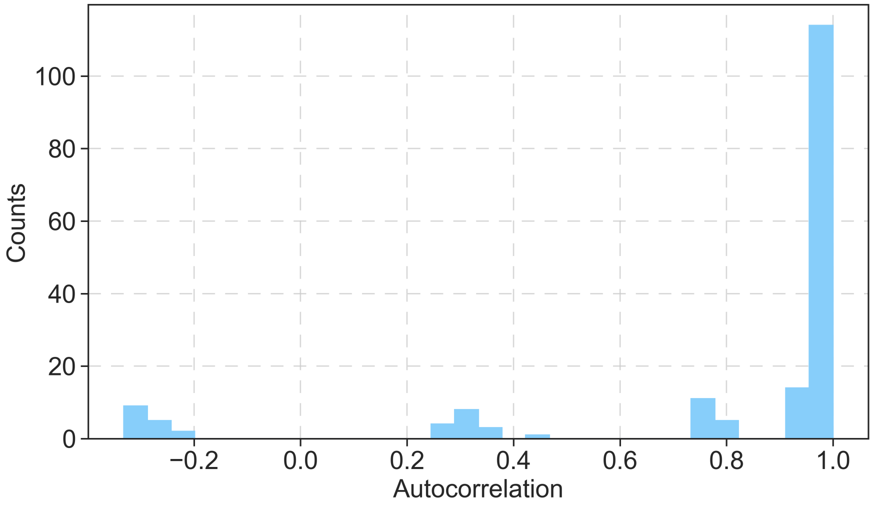


**FIG. S4:** Illustrative example of the signal's autocorrelation distribution across the components produced by the EEMD procedure. The reported distribution refers to the cello in Piece 1, in *let-go* mode.

### S3. NETWORK MEASURES

Here, we provide a brief description of the network measures employed in Sec. II C and Fig. S5. We first introduce the network-based estimators used for the analysis here, then present additional results that complement those in the main text.

#### A. Topological measures

**Clustering and Transitivity**
Clustering, also known as the (local) clustering coefficient (Opsahl and Panzarasa 2009; Watts and Strogatz 1998), is defined, for each node, as the ratio of the number of triangles (three nodes that are all connected) that include the node to the number of possible triangles among its neighbours, and is then averaged across nodes; its weighted version accounts for edge weights. On the other hand, transitivity, or global clustering coefficient, is defined as the ratio of the number of triangles to the number of connected triplets (Luce and Perry 1949; Wasserman and Faust 1994). Both clustering and transitivity aim to describe the likelihood that nodes that share a common neighbour are directly connected, or in other words, the degree to which nodes in a network tend to cluster together. Larger values of these metrics indicate a high level of local interconnectedness, in which neighbours of a node are likely to be neighbours of each other. They can also provide insights into the robustness and redundancy of networks, influencing their resilience to disruptions.

**Square Clustering**
The (average) square clustering coefficient extends the notion of clustering from triangles to squares, i.e. cycles of four nodes (Lind et al. 2005). For each node, it is defined as the fraction of possible squares involving the node and two of its neighbours that are actually present in the network, and it is then averaged across all nodes. This measure is particularly informative in sparse networks, or in networks with few triangles, where the standard clustering coefficient may vanish even in the presence of substantial local cohesion. As for clustering and transitivity, larger values indicate a higher level of local interconnectedness.

**Graph Heterogeneity and Degree Heterogeneity**
Graph heterogeneity and degree heterogeneity both refer to the variability in the structure of a network, encompassing the diversity in the degrees of the nodes (Cohen and Havlin 2010). In particular, graph heterogeneity focuses on the differences in degrees between pairs of connected nodes, while degree heterogeneity is based on the proportions of unique degree values in the entire degree distribution. In a network with high heterogeneity, some nodes have many connections (hubs), while most nodes have few connections, making it more likely that some nodes play critical roles in maintaining connectivity and facilitating the flow of information or resources.

#### Spectral Radius
The spectral radius of a network is the largest absolute eigenvalue of its adjacency matrix. It is a key indicator of various network properties, including stability, connectivity, and dynamic processes such as synchronisation and diffusion (Chung 1997). A larger spectral radius generally indicates a more strongly connected network with more robust structural properties. In practical applications, the spectral radius can help assess the resilience of networks to perturbations and their capacity to support efficient communication or spreading phenomena (Barabási 2013; Boccaletti et al. 2006).

#### Modularity
Modularity is a measure used to quantify the strength of the division of a network into modules or communities, helping to identify functional groups within the network, as well as to understand the organisation and dynamics of complex systems (Newman 2006). High modularity indicates a well-defined community structure, where nodes within the same community have many connections to each other but few to nodes outside the community.

#### Edge Density
Edge density, also known as network density, is a measure of how densely the edges in a network are distributed. It is defined as the ratio of the number of edges in the network to the maximum possible number of edges that could exist between the nodes (Wasserman and Faust 1994). This quantity provides insights into the overall connectivity of the network: a high edge density value indicates a network where nodes are more interconnected, whereas a low one suggests a sparser network (Newman 2018; Rubinov and Sporns 2010).

The results for all eight measures are shown in Fig. S5, and the corresponding tests in Tab. S4. Besides the measures discussed in the main text, weighted clustering is significantly higher in *let-go* in Piece 1 ($t(110) = 2.37$, $p_{\mathrm{FDR}} = 0.036$, $d_z = 0.22$), and transitivity and square clustering in Piece 2 ($t(37) = 2.55$ and 2.68, $p_{\mathrm{FDR}} = 0.030$ for both, $d_z = 0.41$ and 0.44). Graph heterogeneity, for which we had no directional hypothesis, is significantly lower in *let-go* in Piece 1 ($t(110) = -3.15$, $p_{\mathrm{FDR}} = 0.017$, $d_z = -0.30$). Since all 40 tests are corrected together, the spectral radius in Piece 2, the edge density in Piece 4, and the degree heterogeneity in Piece 5, which are significant in Fig. 4b, narrowly miss significance here ($p_{\mathrm{FDR}}$ between 0.054 and 0.065).

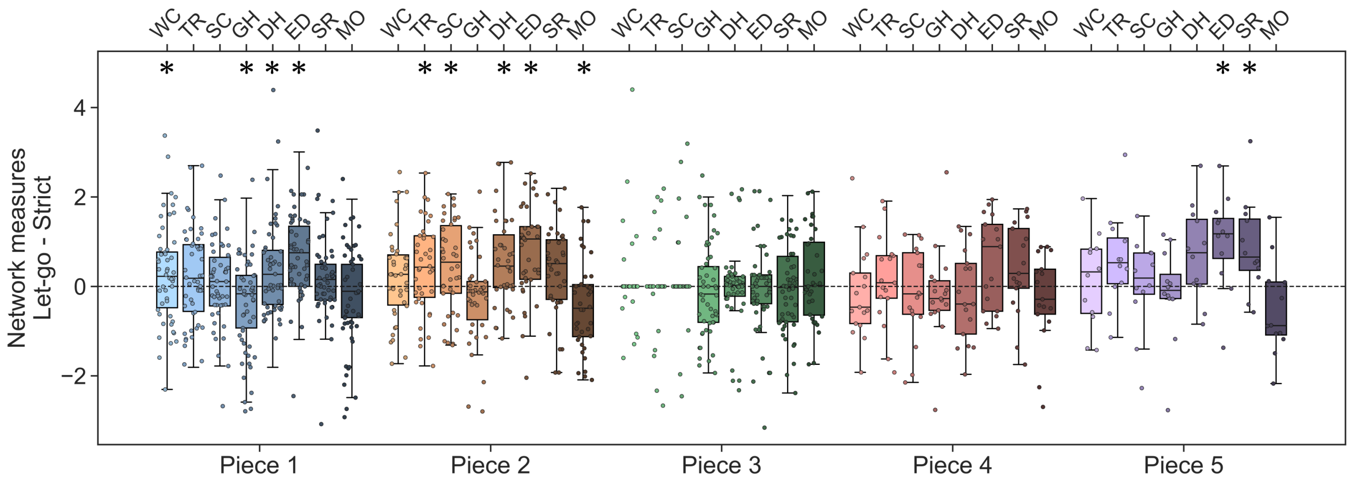


**FIG. S5:** Full list of network measures across all pieces. WC: weighted clustering, TR: transitivity, SC: square clustering, GH: graph heterogeneity, DH: degree heterogeneity, ED: edge density, SR: spectral radius, MO: modularity. Stars mark significant differences (one-sample t-test on the paired differences, FDR-corrected $p < 0.05$ across all 40 tests shown; see Sec. IV G and Tab. S4).

### B. Infomap clusters

In the main text, we showed that cluster entropy is lower in the *let-go* mode, indicating a more cohesive topological structure in this condition. Here, we further support these results by observing that the *let-go* mode is characterised by fewer but larger clusters compared to the *strict* mode (Fig. S6). Both differences are significant in Pieces 1 and 4 (Tab. S3), with the same trend in Pieces 2 and 5, whereas Piece 3 shows no significant difference, consistent with the results of the main text.

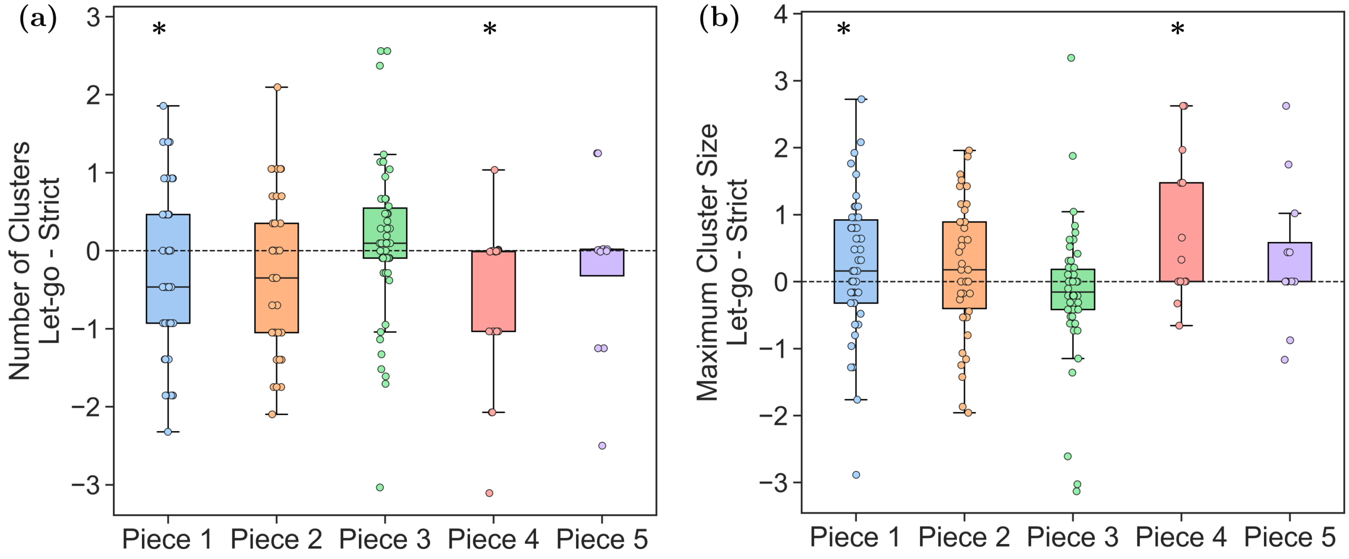


**FIG. S6:** *Let-go–Strict* differences in (a) the number of clusters and (b) the maximum cluster size of PMIME networks. Stars mark significant differences (one-sample t-test on the paired differences, FDR-corrected $p < 0.05$; see Sec. IV G and Tab. S3).

#### C. Normalised PND atoms

In the main text, we showed that *let-go* performances are characterised by more synergistic paths obtained by combining the musicians' brain networks (Fig. 5b). Since *let-go* networks are also denser (Fig. 4b), part of this increase could in principle reflect the larger number of connections rather than a change in how the two brains' networks combine to support communication. To control for this, we normalised the PND atom counts of each epoch by the total number of paths, and repeated the *let-go–strict* comparison (Fig. S7 and Tab. S5).

The results closely match those obtained with the raw counts. Synergy remains significantly higher in *let-go* for Piece 1 ($t(110) = 6.47$, $p_{\mathrm{FDR}} < 10^{-7}$, $d_z = 0.61$), Piece 2 ($t(37) = 3.00$, $p_{\mathrm{FDR}} = 0.022$, $d_z = 0.49$), and Piece 5 ($t(11) = 3.13$, $p_{\mathrm{FDR}} = 0.029$, $d_z = 0.90$), with effect sizes almost identical to the raw ones, and the same positive trend in Piece 4. Unique atoms show no significant difference in any piece. In Piece 3, synergistic paths are almost absent in both conditions, and the normalised unique atoms are widely spread, since the very sparse networks of this dataset contain only a few paths in many epochs (epochs in which the normalisation is undefined are excluded, see Tab. S5). Hence, these results confirm that the increase in synergy during *let-go* is not a mere consequence of the higher number of connections, but reflects a genuine topological feature of communication in the musicians' brains during *let-go* performances.

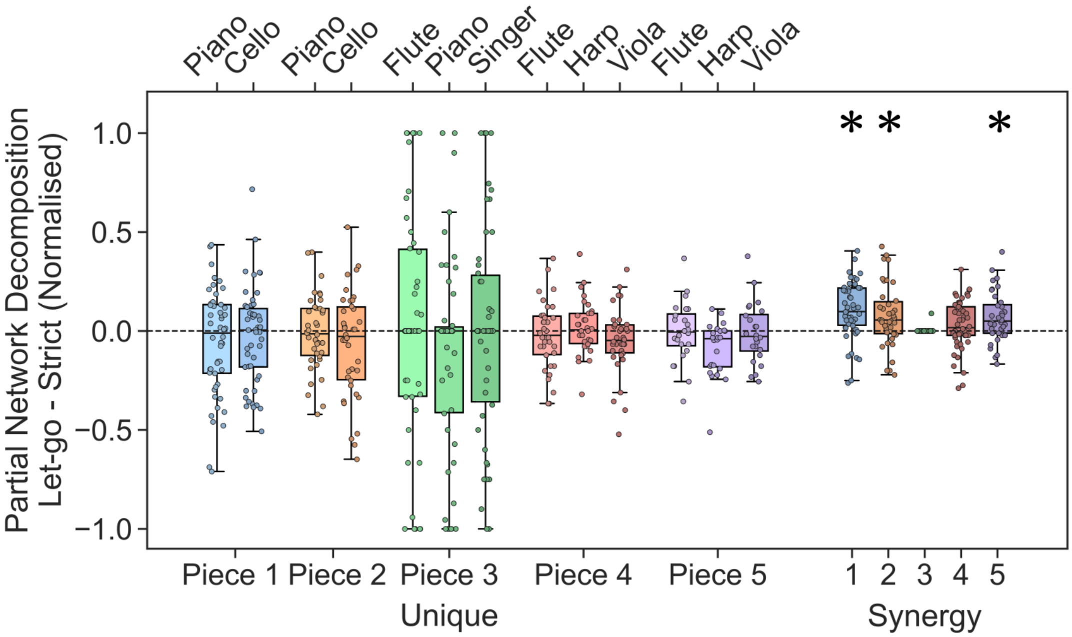


**FIG. S7:** *Let-go–Strict* difference in PND atom counts normalised by the total number of paths. For the trios, PND is computed on all pairs of musicians, and the results are pooled. Stars mark significant differences (one-sample t-test on the paired differences, FDR-corrected $p < 0.05$; see Sec. IV G and Tab. S5).

### S4. STATISTICAL ROBUSTNESS AND FULL RESULTS

To verify that our conclusions do not depend on the parametric assumptions of the t-test, we repeated every statistical test of the main text as a sign-flip permutation test (Maris and Oostenveld 2007; Nichols and Holmes 2002). Under the null hypothesis of no difference between conditions, the *let-go* and *strict* labels within each pair are exchangeable, and swapping them is equivalent to flipping the sign of the corresponding paired difference. We therefore built the null distribution of the mean paired difference by randomly flipping the signs of the differences. For small samples ($n \leq 14$), we enumerated all $2^n$ sign configurations, yielding an exact p-value; otherwise, we drew $N = 100{,}000$ random configurations and computed the p-value as $(k+1)/(N+1)$, where $k$ is the number of permuted statistics at least as extreme as the observed one. Beyond the symmetry of the differences under the null hypothesis, this test makes no assumption on their distribution. We used the same alternative hypotheses (one- or two-sided) and FDR correction as in the main analysis (Sec. IV G).

Results show that the two procedures agree closely: across the 169 tests reported in the figures, the median absolute difference between the uncorrected t-test and permutation p-values is 0.001. After FDR correction, the outcome changes for only three tests, all lying at the significance threshold. Two are significant with the t-test but not with the permutation test: the spectral radius in Piece 2 ($p_{\mathrm{FDR}} = 0.048$ with the t-test vs $p_{\mathrm{perm,FDR}} = 0.051$ with the permutation test) and the edge density in Piece 4 ($p_{\mathrm{FDR}} = 0.049$ vs $p_{\mathrm{perm,FDR}} = 0.054$), both in Fig. 4b. Conversely, the degree heterogeneity in Piece 5 is significant with the permutation test only, within the full set of network measures of Fig. S5 ($p_{\mathrm{FDR}} = 0.054$ vs $p_{\mathrm{perm,FDR}} = 0.046$).

Since epochs are consecutive segments of a continuous recording, we also checked for serial dependence in the epoch-wise paired differences. Their lag-1 autocorrelation is close to zero (median 0.01 across all tests), supporting the treatment of epochs as independent observations.

The complete results of both procedures, including sample sizes, test statistics, and effect sizes, are reported in Tabs. S1–S5.

| Piece | Measure | Alt. | $n$ | $t$ | $d_z$ | $p$ | $p_{\mathrm{FDR}}$ | $p_{\mathrm{perm}}$ | $p_{\mathrm{perm,FDR}}$ |
|---|---|---|---|---|---|---|---|---|---|
| *Broadband CSER (Fig. 2a): channels as samples* | | | | | | | | | |
| 1 | Cello | $>$ | 16 | $-0.23$ | $-0.06$ | 0.588 | 0.588 | 0.596 | 0.596 |
| 1 | Piano | $>$ | 16 | 9.80 | 2.45 | $3.3 \times 10^{-8}$ | $\mathbf{4.2 \times 10^{-7}}$ | $2.0 \times 10^{-5}$ | $\mathbf{2.6 \times 10^{-4}}$ |
| 2 | Cello | $>$ | 16 | 1.53 | 0.38 | 0.073 | 0.111 | 0.073 | 0.123 |
| 2 | Piano | $>$ | 16 | 4.76 | 1.19 | $1.3 \times 10^{-4}$ | $\mathbf{5.5 \times 10^{-4}}$ | $1.2 \times 10^{-4}$ | $\mathbf{7.8 \times 10^{-4}}$ |
| 3 | Flute | $>$ | 18 | 2.34 | 0.55 | 0.016 | **0.041** | 0.017 | **0.044** |
| 3 | Piano | $>$ | 12 | 1.53 | 0.44 | 0.077 | 0.111 | 0.076 | 0.123 |
| 3 | Singer | $>$ | 16 | 1.14 | 0.28 | 0.136 | 0.177 | 0.140 | 0.182 |
| 4 | Flute | $>$ | 10 | 2.17 | 0.69 | 0.029 | 0.063 | 0.023 | 0.051 |
| 4 | Harp | $>$ | 10 | 0.87 | 0.28 | 0.203 | 0.240 | 0.202 | 0.239 |
| 4 | Viola | $>$ | 10 | 1.66 | 0.53 | 0.065 | 0.111 | 0.087 | 0.126 |
| 5 | Flute | $>$ | 10 | 3.59 | 1.13 | 0.003 | **0.010** | $9.8 \times 10^{-4}$ | **0.003** |
| 5 | Harp | $>$ | 10 | 0.58 | 0.18 | 0.288 | 0.312 | 0.291 | 0.315 |
| 5 | Viola | $>$ | 10 | 7.35 | 2.33 | $2.2 \times 10^{-5}$ | $\mathbf{1.4 \times 10^{-4}}$ | $9.8 \times 10^{-4}$ | **0.003** |
| *Broadband CSER (not shown in Fig. 2): channel-averaged values, musician–piece combinations as samples* | | | | | | | | | |
| All | Broadband CSER | $>$ | 13 | 4.61 | 1.28 | $3.0 \times 10^{-4}$ | $\mathbf{3.0 \times 10^{-4}}$ | $2.4 \times 10^{-4}$ | $\mathbf{2.4 \times 10^{-4}}$ |
| *Band-limited CSER (Fig. 2b): musician–piece combinations as samples* | | | | | | | | | |
| All | $\delta$ | $\neq$ | 13 | $-2.39$ | $-0.66$ | 0.034 | **0.043** | 0.037 | **0.046** |
| All | $\theta$ | $\neq$ | 13 | $-3.48$ | $-0.96$ | 0.005 | **0.011** | 0.006 | **0.014** |
| All | $\alpha$ | $\neq$ | 13 | $-2.47$ | $-0.68$ | 0.030 | **0.043** | 0.021 | **0.035** |
| All | $\beta$ | $\neq$ | 13 | $-1.28$ | $-0.35$ | 0.225 | 0.225 | 0.225 | 0.225 |
| All | $\gamma$ | $\neq$ | 13 | 4.39 | 1.22 | $8.8 \times 10^{-4}$ | **0.004** | $9.8 \times 10^{-4}$ | **0.005** |

**TAB. S1:** Statistical tests for the CSER analysis (Fig. 2). The across-musician test of broadband CSER is a single test, and its p-values are therefore not corrected. Alt.: alternative hypothesis for the *let-go*−*strict* difference ($\neq$: two-sided). $n$: number of paired samples; $t$: one-sample t-statistic ($n-1$ degrees of freedom); $d_z$: mean paired difference divided by its standard deviation. $p$ and $p_{\mathrm{perm}}$: uncorrected p-values of the t-test and of the sign-flip permutation test, respectively; $p_{\mathrm{FDR}}$ and $p_{\mathrm{perm,FDR}}$: Benjamini–Hochberg corrected within each figure panel. Corrected p-values below 0.05 are in bold.

| Piece | Measure | Alt. | $n$ | $t$ | $d_z$ | $p$ | $p_{\mathrm{FDR}}$ | $p_{\mathrm{perm}}$ | $p_{\mathrm{perm,FDR}}$ |
|---|---|---|---|---|---|---|---|---|---|
| *Let-go vs strict (Fig. 3a): epochs as samples* | | | | | | | | | |
| 1 | S intra | > | 111 | 2.22 | 0.21 | 0.014 | **0.035** | 0.014 | **0.031** |
| 1 | S inter | > | 111 | 2.84 | 0.27 | 0.003 | **0.014** | 0.002 | **0.011** |
| 1 | O intra | ≠ | 111 | −0.16 | −0.01 | 0.875 | 0.972 | 0.880 | 0.978 |
| 1 | O inter | ≠ | 111 | 1.17 | 0.11 | 0.245 | 0.446 | 0.257 | 0.467 |
| 2 | S intra | > | 38 | 4.87 | 0.79 | $1.0 \times 10^{-5}$ | $\mathbf{2.1 \times 10^{-4}}$ | $1.0 \times 10^{-5}$ | $\mathbf{2.0 \times 10^{-4}}$ |
| 2 | S inter | > | 38 | 4.04 | 0.65 | $1.3 \times 10^{-4}$ | **0.001** | $2.0 \times 10^{-5}$ | $\mathbf{2.0 \times 10^{-4}}$ |
| 2 | O intra | ≠ | 38 | −0.23 | −0.04 | 0.819 | 0.972 | 0.820 | 0.978 |
| 2 | O inter | ≠ | 38 | 1.86 | 0.30 | 0.071 | 0.143 | 0.066 | 0.131 |
| 3 | S intra | > | 60 | −2.86 | −0.37 | 0.997 | 0.997 | 0.997 | 0.997 |
| 3 | S inter | > | 60 | −2.39 | −0.31 | 0.990 | 0.997 | 0.990 | 0.997 |
| 3 | O intra | ≠ | 60 | −2.49 | −0.32 | 0.016 | **0.035** | 0.014 | **0.031** |
| 3 | O inter | ≠ | 60 | −0.20 | −0.03 | 0.842 | 0.972 | 0.846 | 0.978 |
| 4 | S intra | > | 17 | 2.75 | 0.67 | 0.007 | **0.024** | 0.006 | **0.019** |
| 4 | S inter | > | 17 | 2.40 | 0.58 | 0.015 | **0.035** | 0.011 | **0.031** |
| 4 | O intra | ≠ | 17 | −0.32 | −0.08 | 0.752 | 0.972 | 0.783 | 0.978 |
| 4 | O inter | ≠ | 17 | 1.04 | 0.25 | 0.313 | 0.522 | 0.390 | 0.651 |
| 5 | S intra | > | 12 | 3.29 | 0.95 | 0.004 | **0.014** | 0.003 | **0.011** |
| 5 | S inter | > | 12 | 4.46 | 1.29 | $4.8 \times 10^{-4}$ | **0.003** | $2.4 \times 10^{-4}$ | **0.002** |
| 5 | O intra | ≠ | 12 | 0.39 | 0.11 | 0.706 | 0.972 | 0.719 | 0.978 |
| 5 | O inter | ≠ | 12 | −0.55 | −0.16 | 0.591 | 0.910 | 0.638 | 0.978 |
| *Intra- vs inter-brain triplets (Fig. 3b): epochs as samples* | | | | | | | | | |
| 1 | S, intra vs inter (*strict*) | > | 112 | 15.05 | 1.42 | $7.1 \times 10^{-29}$ | $\mathbf{3.6 \times 10^{-28}}$ | $1.0 \times 10^{-5}$ | $\mathbf{2.9 \times 10^{-5}}$ |
| 1 | S, intra vs inter (*let-go*) | > | 111 | 17.30 | 1.64 | $1.9 \times 10^{-33}$ | $\mathbf{1.9 \times 10^{-32}}$ | $1.0 \times 10^{-5}$ | $\mathbf{2.9 \times 10^{-5}}$ |
| 1 | O, intra vs inter (*strict*) | ≠ | 112 | 1.43 | 0.13 | 0.157 | 0.241 | 0.183 | 0.261 |
| 1 | O, intra vs inter (*let-go*) | ≠ | 111 | 1.22 | 0.12 | 0.226 | 0.301 | 0.240 | 0.320 |
| 2 | S, intra vs inter (*strict*) | > | 38 | 10.37 | 1.68 | $8.5 \times 10^{-13}$ | $\mathbf{3.4 \times 10^{-12}}$ | $1.0 \times 10^{-5}$ | $\mathbf{2.9 \times 10^{-5}}$ |
| 2 | S, intra vs inter (*let-go*) | > | 38 | 10.18 | 1.65 | $1.4 \times 10^{-12}$ | $\mathbf{4.7 \times 10^{-12}}$ | $1.0 \times 10^{-5}$ | $\mathbf{2.9 \times 10^{-5}}$ |
| 2 | O, intra vs inter (*strict*) | ≠ | 38 | 0.61 | 0.10 | 0.546 | 0.642 | 0.549 | 0.646 |
| 2 | O, intra vs inter (*let-go*) | ≠ | 38 | −1.16 | −0.19 | 0.254 | 0.318 | 0.258 | 0.322 |
| 3 | S, intra vs inter (*strict*) | > | 60 | 25.69 | 3.32 | $5.0 \times 10^{-34}$ | $\mathbf{1.0 \times 10^{-32}}$ | $1.0 \times 10^{-5}$ | $\mathbf{2.9 \times 10^{-5}}$ |
| 3 | S, intra vs inter (*let-go*) | > | 61 | 20.75 | 2.66 | $2.3 \times 10^{-29}$ | $\mathbf{1.5 \times 10^{-28}}$ | $1.0 \times 10^{-5}$ | $\mathbf{2.9 \times 10^{-5}}$ |
| 3 | O, intra vs inter (*strict*) | ≠ | 60 | 1.39 | 0.18 | 0.170 | 0.243 | 0.173 | 0.261 |
| 3 | O, intra vs inter (*let-go*) | ≠ | 61 | −2.59 | −0.33 | 0.012 | **0.022** | 0.013 | **0.023** |
| 4 | S, intra vs inter (*strict*) | > | 17 | 5.26 | 1.28 | $3.9 \times 10^{-5}$ | $\mathbf{8.7 \times 10^{-5}}$ | $2.0 \times 10^{-5}$ | $\mathbf{5.0 \times 10^{-5}}$ |
| 4 | S, intra vs inter (*let-go*) | > | 17 | 5.29 | 1.28 | $3.7 \times 10^{-5}$ | $\mathbf{8.7 \times 10^{-5}}$ | $1.0 \times 10^{-5}$ | $\mathbf{2.9 \times 10^{-5}}$ |
| 4 | O, intra vs inter (*strict*) | ≠ | 17 | 1.92 | 0.47 | 0.072 | 0.121 | 0.071 | 0.119 |
| 4 | O, intra vs inter (*let-go*) | ≠ | 17 | 0.37 | 0.09 | 0.718 | 0.755 | 0.819 | 0.862 |
| 5 | S, intra vs inter (*strict*) | > | 12 | 5.39 | 1.56 | $1.1 \times 10^{-4}$ | $\mathbf{2.2 \times 10^{-4}}$ | $2.4 \times 10^{-4}$ | $\mathbf{4.9 \times 10^{-4}}$ |
| 5 | S, intra vs inter (*let-go*) | > | 13 | 7.48 | 2.07 | $3.7 \times 10^{-6}$ | $\mathbf{1.1 \times 10^{-5}}$ | $1.2 \times 10^{-4}$ | $\mathbf{2.7 \times 10^{-4}}$ |
| 5 | O, intra vs inter (*strict*) | ≠ | 12 | −0.44 | −0.13 | 0.667 | 0.741 | 0.667 | 0.742 |
| 5 | O, intra vs inter (*let-go*) | ≠ | 13 | 0.04 | 0.01 | 0.969 | 0.969 | 0.968 | 0.968 |

**TAB. S2:** Statistical tests for the S- and O-information analysis (Fig. 3). Alt.: alternative hypothesis for the *let-go*−*strict* difference (≠: two-sided); for the intra- vs inter-brain comparison, Alt. refers to the intra−inter difference. $n$: number of paired samples; $t$: one-sample t-statistic ($n-1$ degrees of freedom); $d_z$: mean paired difference divided by its standard deviation. $p$ and $p_{\mathrm{perm}}$: uncorrected p-values of the t-test and of the sign-flip permutation test, respectively; $p_{\mathrm{FDR}}$ and $p_{\mathrm{perm,FDR}}$: Benjamini–Hochberg corrected within each figure panel. Corrected p-values below 0.05 are in bold.

| Piece | Measure | Alt. | $n$ | $t$ | $d_z$ | $p$ | $p_{\mathrm{FDR}}$ | $p_{\mathrm{perm}}$ | $p_{\mathrm{perm,FDR}}$ |
|---|---|---|---|---|---|---|---|---|---|
| *Cluster entropy (Fig. 4a)* | | | | | | | | | |
| 1 | Cluster entropy | < | 111 | −3.82 | −0.36 | $1.1 \times 10^{-4}$ | $\mathbf{5.5 \times 10^{-4}}$ | $1.4 \times 10^{-4}$ | $\mathbf{7.0 \times 10^{-4}}$ |
| 2 | Cluster entropy | < | 38 | −1.48 | −0.24 | 0.073 | 0.122 | 0.071 | 0.119 |
| 3 | Cluster entropy | < | 60 | −1.02 | −0.13 | 0.157 | 0.173 | 0.160 | 0.180 |
| 4 | Cluster entropy | < | 17 | −2.64 | −0.64 | 0.009 | **0.022** | 0.010 | **0.026** |
| 5 | Cluster entropy | < | 12 | −0.98 | −0.28 | 0.173 | 0.173 | 0.180 | 0.180 |
| *Network measures (Fig. 4b)* | | | | | | | | | |
| 1 | Degree heterogeneity | > | 111 | 3.06 | 0.29 | 0.001 | **0.007** | 0.001 | **0.006** |
| 1 | Edge density | > | 111 | 6.73 | 0.64 | $4.0 \times 10^{-10}$ | $\mathbf{7.9 \times 10^{-9}}$ | $1.0 \times 10^{-5}$ | $\mathbf{2.0 \times 10^{-4}}$ |
| 1 | Modularity | < | 111 | −1.31 | −0.12 | 0.096 | 0.148 | 0.095 | 0.146 |
| 1 | Spectral radius | > | 111 | 0.47 | 0.05 | 0.318 | 0.424 | 0.319 | 0.426 |
| 2 | Degree heterogeneity | > | 38 | 3.46 | 0.56 | $6.8 \times 10^{-4}$ | **0.005** | $6.2 \times 10^{-4}$ | **0.004** |
| 2 | Edge density | > | 38 | 4.76 | 0.77 | $1.5 \times 10^{-5}$ | $\mathbf{1.5 \times 10^{-4}}$ | $6.0 \times 10^{-5}$ | $\mathbf{6.0 \times 10^{-4}}$ |
| 2 | Modularity | < | 38 | −2.58 | −0.42 | 0.007 | **0.020** | 0.007 | **0.021** |
| 2 | Spectral radius | > | 38 | 2.09 | 0.34 | 0.022 | **0.048** | 0.023 | 0.051 |
| 3 | Degree heterogeneity | > | 60 | −0.92 | −0.12 | 0.818 | 0.861 | 0.817 | 0.860 |
| 3 | Edge density | > | 60 | 0.02 | 0.00 | 0.490 | 0.613 | 0.491 | 0.614 |
| 3 | Modularity | < | 36 | 1.13 | 0.19 | 0.866 | 0.866 | 0.866 | 0.866 |
| 3 | Spectral radius | > | 60 | −0.78 | −0.10 | 0.780 | 0.861 | 0.782 | 0.860 |
| 4 | Degree heterogeneity | > | 17 | −0.81 | −0.20 | 0.784 | 0.861 | 0.786 | 0.860 |
| 4 | Edge density | > | 17 | 2.13 | 0.52 | 0.025 | **0.049** | 0.027 | 0.054 |
| 4 | Modularity | < | 17 | −1.07 | −0.26 | 0.151 | 0.215 | 0.158 | 0.226 |
| 4 | Spectral radius | > | 17 | 1.59 | 0.38 | 0.066 | 0.110 | 0.066 | 0.109 |
| 5 | Degree heterogeneity | > | 12 | 2.44 | 0.70 | 0.016 | **0.041** | 0.014 | **0.034** |
| 5 | Edge density | > | 12 | 3.40 | 0.98 | 0.003 | **0.012** | 0.005 | **0.016** |
| 5 | Modularity | < | 12 | −1.75 | −0.51 | 0.054 | 0.098 | 0.055 | 0.100 |
| 5 | Spectral radius | > | 12 | 2.93 | 0.85 | 0.007 | **0.020** | 0.005 | **0.016** |
| *Number of clusters (Fig. S6a)* | | | | | | | | | |
| 1 | Number of clusters | < | 111 | −5.36 | −0.51 | $2.4 \times 10^{-7}$ | $\mathbf{1.2 \times 10^{-6}}$ | $1.0 \times 10^{-5}$ | $\mathbf{5.0 \times 10^{-5}}$ |
| 2 | Number of clusters | < | 38 | −1.79 | −0.29 | 0.041 | 0.068 | 0.044 | 0.073 |
| 3 | Number of clusters | < | 60 | 0.81 | 0.10 | 0.790 | 0.790 | 0.792 | 0.792 |
| 4 | Number of clusters | < | 17 | −2.68 | −0.65 | 0.008 | **0.021** | 0.015 | **0.036** |
| 5 | Number of clusters | < | 12 | −0.69 | −0.20 | 0.252 | 0.315 | 0.375 | 0.469 |
| *Maximum cluster size (Fig. S6b)* | | | | | | | | | |
| 1 | Max cluster size | > | 111 | 2.30 | 0.22 | 0.012 | **0.037** | 0.012 | **0.050** |
| 2 | Max cluster size | > | 38 | 1.20 | 0.19 | 0.119 | 0.167 | 0.123 | 0.186 |
| 3 | Max cluster size | > | 60 | −1.05 | −0.14 | 0.851 | 0.851 | 0.849 | 0.849 |
| 4 | Max cluster size | > | 17 | 2.39 | 0.58 | 0.015 | **0.037** | 0.020 | **0.050** |
| 5 | Max cluster size | > | 12 | 1.17 | 0.34 | 0.134 | 0.167 | 0.148 | 0.186 |

**TAB. S3:** Statistical tests for the analysis of PMIME networks (Figs. 4 and S6); epochs as samples. Modularity is undefined for networks with fewer than 10 edges, and the corresponding epoch pairs are excluded. Alt.: alternative hypothesis for the *let-go*−*strict* difference ($\neq$: two-sided). $n$: number of paired samples; $t$: one-sample t-statistic ($n-1$ degrees of freedom); $d_z$: mean paired difference divided by its standard deviation. $p$ and $p_{\mathrm{perm}}$: uncorrected p-values of the t-test and of the sign-flip permutation test, respectively; $p_{\mathrm{FDR}}$ and $p_{\mathrm{perm,FDR}}$: Benjamini–Hochberg corrected within each figure panel. Corrected p-values below 0.05 are in bold.

| Piece | Measure | Alt. | $n$ | $t$ | $d_z$ | $p$ | $p_{\mathrm{FDR}}$ | $p_{\mathrm{perm}}$ | $p_{\mathrm{perm,FDR}}$ |
|---|---|---|---|---|---|---|---|---|---|
| *All network measures (Fig. S5): one FDR family of 40 tests* | | | | | | | | | |
| 1 | Weighted clustering | > | 111 | 2.37 | 0.22 | 0.010 | **0.036** | 0.009 | **0.034** |
| 1 | Transitivity | > | 111 | 2.14 | 0.20 | 0.017 | 0.054 | 0.017 | 0.051 |
| 1 | Square clustering | > | 111 | 0.21 | 0.02 | 0.418 | 0.582 | 0.418 | 0.582 |
| 1 | Graph heterogeneity | $\neq$ | 111 | −3.15 | −0.30 | 0.002 | **0.017** | 0.002 | **0.017** |
| 1 | Degree heterogeneity | > | 111 | 3.06 | 0.29 | 0.001 | **0.014** | 0.001 | **0.012** |
| 1 | Edge density | > | 111 | 6.73 | 0.64 | $4.0 \times 10^{-10}$ | $\mathbf{1.6 \times 10^{-8}}$ | $1.0 \times 10^{-5}$ | $\mathbf{4.0 \times 10^{-4}}$ |
| 1 | Modularity | < | 111 | −1.31 | −0.12 | 0.096 | 0.193 | 0.097 | 0.194 |
| 1 | Spectral radius | > | 111 | 0.47 | 0.05 | 0.318 | 0.482 | 0.320 | 0.481 |
| 2 | Weighted clustering | > | 38 | 1.33 | 0.22 | 0.096 | 0.193 | 0.096 | 0.194 |
| 2 | Transitivity | > | 38 | 2.55 | 0.41 | 0.007 | **0.030** | 0.008 | **0.033** |
| 2 | Square clustering | > | 38 | 2.68 | 0.44 | 0.005 | **0.030** | 0.005 | **0.027** |
| 2 | Graph heterogeneity | $\neq$ | 38 | −1.57 | −0.26 | 0.124 | 0.236 | 0.125 | 0.238 |
| 2 | Degree heterogeneity | > | 38 | 3.46 | 0.56 | $6.8 \times 10^{-4}$ | **0.009** | $5.2 \times 10^{-4}$ | **0.007** |
| 2 | Edge density | > | 38 | 4.76 | 0.77 | $1.5 \times 10^{-5}$ | $\mathbf{2.9 \times 10^{-4}}$ | $5.0 \times 10^{-5}$ | **0.001** |
| 2 | Modularity | < | 38 | −2.58 | −0.42 | 0.007 | **0.030** | 0.007 | **0.032** |
| 2 | Spectral radius | > | 38 | 2.09 | 0.34 | 0.022 | 0.062 | 0.023 | 0.064 |
| 3 | Weighted clustering | > | 36 | 0.78 | 0.13 | 0.221 | 0.368 | 0.240 | 0.399 |
| 3 | Transitivity | > | 36 | 0.16 | 0.03 | 0.437 | 0.582 | 0.436 | 0.582 |
| 3 | Square clustering | > | 36 | 1.00 | 0.17 | 0.163 | 0.284 | 0.171 | 0.298 |
| 3 | Graph heterogeneity | $\neq$ | 60 | −0.63 | −0.08 | 0.532 | 0.645 | 0.533 | 0.646 |
| 3 | Degree heterogeneity | > | 60 | −0.92 | −0.12 | 0.818 | 0.839 | 0.819 | 0.840 |
| 3 | Edge density | > | 60 | 0.02 | 0.00 | 0.490 | 0.632 | 0.496 | 0.637 |
| 3 | Modularity | < | 36 | 1.13 | 0.19 | 0.866 | 0.866 | 0.867 | 0.867 |
| 3 | Spectral radius | > | 60 | −0.78 | −0.10 | 0.780 | 0.825 | 0.782 | 0.829 |
| 4 | Weighted clustering | > | 17 | −0.59 | −0.14 | 0.717 | 0.797 | 0.716 | 0.797 |
| 4 | Transitivity | > | 17 | 0.46 | 0.11 | 0.325 | 0.482 | 0.325 | 0.481 |
| 4 | Square clustering | > | 17 | −0.58 | −0.14 | 0.715 | 0.797 | 0.717 | 0.797 |
| 4 | Graph heterogeneity | $\neq$ | 17 | −0.68 | −0.16 | 0.506 | 0.633 | 0.510 | 0.637 |
| 4 | Degree heterogeneity | > | 17 | −0.81 | −0.20 | 0.784 | 0.825 | 0.788 | 0.829 |
| 4 | Edge density | > | 17 | 2.13 | 0.52 | 0.025 | 0.065 | 0.026 | 0.069 |
| 4 | Modularity | < | 17 | −1.07 | −0.26 | 0.151 | 0.274 | 0.155 | 0.281 |
| 4 | Spectral radius | > | 17 | 1.59 | 0.38 | 0.066 | 0.147 | 0.066 | 0.147 |
| 5 | Weighted clustering | > | 12 | 0.51 | 0.15 | 0.310 | 0.482 | 0.308 | 0.481 |
| 5 | Transitivity | > | 12 | 1.95 | 0.56 | 0.039 | 0.097 | 0.037 | 0.093 |
| 5 | Square clustering | > | 12 | 0.18 | 0.05 | 0.430 | 0.582 | 0.429 | 0.582 |
| 5 | Graph heterogeneity | $\neq$ | 12 | −0.49 | −0.14 | 0.636 | 0.748 | 0.688 | 0.797 |
| 5 | Degree heterogeneity | > | 12 | 2.44 | 0.70 | 0.016 | 0.054 | 0.014 | **0.046** |
| 5 | Edge density | > | 12 | 3.40 | 0.98 | 0.003 | **0.020** | 0.005 | **0.027** |
| 5 | Modularity | < | 12 | −1.75 | −0.51 | 0.054 | 0.127 | 0.055 | 0.129 |
| 5 | Spectral radius | > | 12 | 2.93 | 0.85 | 0.007 | **0.030** | 0.005 | **0.027** |

**TAB. S4:** Statistical tests for the full set of network measures (Fig. S5); epochs as samples. Clustering coefficients and modularity are undefined for networks with fewer than 10 edges, and the corresponding epoch pairs are excluded. Alt.: alternative hypothesis for the *let-go−strict* difference ($\neq$: two-sided). $n$: number of paired samples; $t$: one-sample t-statistic ($n-1$ degrees of freedom); $d_z$: mean paired difference divided by its standard deviation. $p$ and $p_{\mathrm{perm}}$: uncorrected p-values of the t-test and of the sign-flip permutation test, respectively; $p_{\mathrm{FDR}}$ and $p_{\mathrm{perm,FDR}}$: Benjamini–Hochberg corrected across all 40 tests. Corrected p-values below 0.05 are in bold.

| Piece | Measure | Alt. | $n$ | $t$ | $d_z$ | $p$ | $p_{\mathrm{FDR}}$ | $p_{\mathrm{perm}}$ | $p_{\mathrm{perm,FDR}}$ |
|---|---|---|---|---|---|---|---|---|---|
| *Raw PND atom counts (Fig. 5b)* | | | | | | | | | |
| 1 | Unique Piano | $\neq$ | 111 | −0.35 | −0.03 | 0.726 | 0.887 | 0.726 | 0.890 |
| 1 | Unique Cello | $\neq$ | 111 | 1.55 | 0.15 | 0.124 | 0.372 | 0.125 | 0.374 |
| 1 | Synergy | > | 111 | 6.60 | 0.63 | $7.5 \times 10^{-10}$ | $\mathbf{1.4 \times 10^{-8}}$ | $1.0 \times 10^{-5}$ | $\mathbf{1.8 \times 10^{-4}}$ |
| 2 | Unique Piano | $\neq$ | 38 | 1.31 | 0.21 | 0.199 | 0.447 | 0.203 | 0.461 |
| 2 | Unique Cello | $\neq$ | 38 | −0.07 | −0.01 | 0.942 | 0.942 | 0.945 | 0.945 |
| 2 | Synergy | > | 38 | 3.14 | 0.51 | 0.002 | **0.015** | 0.002 | **0.014** |
| 3 | Unique Flute | $\neq$ | 60 | 1.11 | 0.14 | 0.271 | 0.487 | 0.286 | 0.571 |
| 3 | Unique Piano | $\neq$ | 60 | −0.18 | −0.02 | 0.862 | 0.912 | 0.871 | 0.922 |
| 3 | Unique Singer | $\neq$ | 60 | −0.97 | −0.12 | 0.338 | 0.507 | 0.401 | 0.656 |
| 3 | Synergy | > | 60 | 0.72 | 0.09 | 0.236 | 0.471 | 0.503 | 0.744 |
| 4 | Unique Flute | $\neq$ | 17 | −0.66 | −0.16 | 0.519 | 0.718 | 0.538 | 0.744 |
| 4 | Unique Harp | $\neq$ | 17 | 1.04 | 0.25 | 0.313 | 0.507 | 0.317 | 0.571 |
| 4 | Unique Viola | $\neq$ | 17 | −1.34 | −0.33 | 0.198 | 0.447 | 0.205 | 0.461 |
| 4 | Synergy | > | 17 | 1.58 | 0.38 | 0.067 | 0.242 | 0.067 | 0.241 |
| 5 | Unique Flute | $\neq$ | 12 | −0.34 | −0.10 | 0.739 | 0.887 | 0.742 | 0.890 |
| 5 | Unique Harp | $\neq$ | 12 | −2.78 | −0.80 | 0.018 | 0.080 | 0.016 | 0.070 |
| 5 | Unique Viola | $\neq$ | 12 | −0.22 | −0.06 | 0.829 | 0.912 | 0.839 | 0.922 |
| 5 | Synergy | > | 12 | 3.17 | 0.91 | 0.004 | **0.027** | 0.005 | **0.031** |
| *Normalised PND atom counts (Fig. S7)* | | | | | | | | | |
| 1 | Unique Piano | $\neq$ | 111 | −2.51 | −0.24 | 0.014 | 0.058 | 0.013 | 0.056 |
| 1 | Unique Cello | $\neq$ | 111 | −1.04 | −0.10 | 0.300 | 0.541 | 0.301 | 0.602 |
| 1 | Synergy | > | 111 | 6.47 | 0.61 | $1.4 \times 10^{-9}$ | $\mathbf{2.5 \times 10^{-8}}$ | $1.0 \times 10^{-5}$ | $\mathbf{1.8 \times 10^{-4}}$ |
| 2 | Unique Piano | $\neq$ | 38 | −0.28 | −0.04 | 0.784 | 0.882 | 0.782 | 0.880 |
| 2 | Unique Cello | $\neq$ | 38 | −1.58 | −0.26 | 0.123 | 0.316 | 0.124 | 0.318 |
| 2 | Synergy | > | 38 | 3.00 | 0.49 | 0.002 | **0.022** | 0.002 | **0.021** |
| 3 | Unique Flute | $\neq$ | 42 | 0.02 | 0.00 | 0.984 | 0.984 | 0.984 | 0.984 |
| 3 | Unique Piano | $\neq$ | 42 | −0.14 | −0.02 | 0.892 | 0.944 | 0.893 | 0.945 |
| 3 | Unique Singer | $\neq$ | 42 | 0.57 | 0.09 | 0.573 | 0.794 | 0.574 | 0.795 |
| 3 | Synergy | > | 42 | 1.00 | 0.15 | 0.162 | 0.364 | 0.500 | 0.749 |
| 4 | Unique Flute | $\neq$ | 17 | −0.78 | −0.19 | 0.449 | 0.673 | 0.462 | 0.749 |
| 4 | Unique Harp | $\neq$ | 17 | 0.95 | 0.23 | 0.355 | 0.582 | 0.351 | 0.632 |
| 4 | Unique Viola | $\neq$ | 17 | −1.39 | −0.34 | 0.184 | 0.368 | 0.187 | 0.421 |
| 4 | Synergy | > | 17 | 1.56 | 0.38 | 0.070 | 0.209 | 0.069 | 0.207 |
| 5 | Unique Flute | $\neq$ | 12 | −0.42 | −0.12 | 0.680 | 0.875 | 0.678 | 0.872 |
| 5 | Unique Harp | $\neq$ | 12 | −2.84 | −0.82 | 0.016 | 0.058 | 0.016 | 0.056 |
| 5 | Unique Viola | $\neq$ | 12 | −0.29 | −0.08 | 0.779 | 0.882 | 0.773 | 0.880 |
| 5 | Synergy | > | 12 | 3.13 | 0.90 | 0.005 | **0.029** | 0.005 | **0.032** |

**TAB. S5:** Statistical tests for the Partial Network Decomposition (Figs. 5 and S7); epochs as samples. For the trios, PND atoms are averaged across pairs of musicians within each epoch. Alt.: alternative hypothesis for the *let-go−strict* difference ($\neq$: two-sided). $n$: number of paired samples; $t$: one-sample t-statistic ($n-1$ degrees of freedom); $d_z$: mean paired difference divided by its standard deviation. $p$ and $p_{\mathrm{perm}}$: uncorrected p-values of the t-test and of the sign-flip permutation test, respectively; $p_{\mathrm{FDR}}$ and $p_{\mathrm{perm,FDR}}$: Benjamini–Hochberg corrected within each figure panel. Corrected p-values below 0.05 are in bold.

Barabási, A.-L. (2013) 'Network science', *Philosophical Transactions of the Royal Society A: Mathematical, Physical and Engineering Sciences*, 371(1987), p. 20120375.

Boccaletti, S., Latora, V., Moreno, Y., Chavez, M. and Hwang, D.-U. (2006) 'Complex networks: Structure and dynamics', *Physics Reports*, 424(4-5), pp. 175–308.

Chen, X., Liu, Q., Tao, W., Li, L., Lee, S., Liu, A., Chen, Q., Cheng, J., McKeown, M. J. and Wang, Z. J. (2019a) 'Remae: User-friendly toolbox for removing muscle artifacts from eeg', *IEEE Transactions on Instrumentation and Measurement*, 69(5), pp. 2105–2119.

Chen, X., Xu, X., Liu, A., Lee, S., Chen, X., Zhang, X., McKeown, M. J. and Wang, Z. J. (2019b) 'Removal of muscle artifacts from the eeg: A review and recommendations', *IEEE Sensors Journal*, 19(14), pp. 5353–5368.

Chung, F. R. (1997) *Spectral graph theory*, vol. 92, American Mathematical Soc.

Cohen, R. and Havlin, S. (2010) *Complex networks: structure, robustness and function*, Cambridge university press.

De Clercq, W., Vergult, A., Vanrumste, B., Van Paesschen, W. and Van Huffel, S. (2006) 'Canonical correlation analysis applied to remove muscle artifacts from the electroencephalogram', *IEEE Transactions on Biomedical Engineering*, 53(12), pp. 2583–2587.

Lind, P. G., González, M. C. and Herrmann, H. J. (2005) 'Cycles and clustering in bipartite networks', *Physical Review E*, 72(5), p. 056127.

Luce, R. D. and Perry, A. D. (1949) 'A method of matrix analysis of group structure', *Psychometrika*, 14(2), pp. 95–116.

Maris, E. and Oostenveld, R. (2007) 'Nonparametric statistical testing of eeg-and meg-data', *Journal of Neuroscience Methods*, 164(1), pp. 177–190.

Newman, M. (2018) *Networks*, Oxford university press.

Newman, M. E. (2006) 'Modularity and community structure in networks', *Proceedings of the National Academy of Sciences*, 103(23), pp. 8577–8582.

Nichols, T. E. and Holmes, A. P. (2002) 'Nonparametric permutation tests for functional neuroimaging: A primer with examples', *Human Brain Mapping*, 15(1), pp. 1–25.

Opsahl, T. and Panzarasa, P. (2009) 'Clustering in weighted networks', *Social Networks*, 31(2), pp. 155–163.

Rubinov, M. and Sporns, O. (2010) 'Complex network measures of brain connectivity: uses and interpretations', *Neuroimage*, 52(3), pp. 1059–1069.

Wasserman, S. and Faust, K. (1994) *Social network analysis: Methods and applications*, Cambridge university press.

Watts, D. J. and Strogatz, S. H. (1998) 'Collective dynamics of 'small-world'networks', *Nature*, 393(6684), pp. 440–442.

Wu, Z. and Huang, N. E. (2009) 'Ensemble empirical mode decomposition: a noise-assisted data analysis method', *Advances in Adaptive Data Analysis*, 1(01), pp. 1–41.
